# The RNA Polymerase II core subunit RPB-9 directs transcriptional elongation at piRNA loci in *Caenorhabditis elegans*

**DOI:** 10.1101/2020.05.01.070433

**Authors:** Ahmet C. Berkyurek, Giulia Furlan, Lisa Lampersberger, Toni Beltran, Eva-Maria Weick, Emily Nischwitz, Isabela Cunha Navarro, Fabian Braukmann, Alper Akay, Jonathan Price, Falk Butter, Peter Sarkies, Eric A. Miska

## Abstract

PIWI-interacting RNAs (piRNAs) are genome-encoded small RNAs that regulate germ cell development and maintain germline integrity in many animals. Mature piRNAs engage Piwi Argonaute proteins to silence complementary transcripts, including transposable elements and endogenous genes. piRNA biogenesis mechanisms are diverse and remain poorly understood. Here, we identify the RNA Polymerase II (RNA Pol II) core subunit RPB-9 as required for piRNA-mediated silencing in the nematode *Caenorhabditis elegans. rpb-9* mutants fail to initiate heritable piRNA-mediated gene silencing. Furthermore, we show that RPB-9 is required to repress two DNA transposon families and a subset of somatic genes in the *C. elegans* germline. We provide genetic and biochemical evidence that RPB-9 is required for piRNA biogenesis. We demonstrate that RPB-9 acts to promote transcriptional elongation/termination at endogenous piRNA loci. We conclude that as a part of its rapid evolution the piRNA pathway has co-opted another ancient machinery, this time for high-fidelity transcription.

## INTRODUCTION

The PIWI-interacting RNA (piRNA) pathway protects animal germlines from transposable elements and other selfish genetic parasites, thereby ensuring transgenerational genomic integrity and fertility (Bagijn et al., 2012a; Lee et al., 2012). The piRNA pathway is widespread in metazoans and relies on the cooperation between small RNAs and Argonaute (Ago) proteins to silence targets both at the post-transcriptional and at the co-transcriptional levels; the precise molecular mechanisms through which it operates however vary between species.

In *C. elegans*, piRNAs play a fundamental role in recognizing non-self DNA elements, inducing their repression and promoting a multigenerational epigenetic memory of their silencing (Ashe et al., 2012; Bagijn et al., 2012a; Lee et al., 2012; Shirayama et al., 2012). Moreover, they have been shown to regulate the expression of a subset of endogenous genes in the germline (Rojas-Ríos and Simonelig, 2018). Most *C. elegans* piRNAs are encoded by two specific clusters on chromosome IV, which contain thousands of intergenic and intronic individual piRNA transcription units enriched in Ruby motif-containing promoters (motif-dependent or type I piRNAs) (Billi et al., 2013; Cecere et al., 2012; Gu et al., 2012; Ruby et al., 2006). A smaller number of piRNAs are produced from other loci in the genome (Batista et al., 2008) and transcribed bidirectionally from transcription start sites (TSSs) flanked by YRNT motifs (motif-independent or type II piRNAs) (Gu et al., 2012; Ruby et al., 2006).

Transcription at motif-dependent piRNA loci requires Forkhead transcription factors (Cecere et al., 2012), and depends on the presence of SNPC-4 (Kasper et al., 2014) and the USTC complex, composed of PRDE-1, SNPC-4, TOFU-4 and TOFU-5 (Weick et al., 2014; Weng et al., 2019) at the Ruby motif. Transcribed piRNA precursors are 26 to 29 nucleotides (nt) in length and carry a 5′ 7-methylguanylate cap (Goh et al., 2014; Gu et al., 2012; Weick et al., 2014). After transcription, precursors are promptly exported out of the nucleus, possibly by PID-1 (de Albuquerque et al., 2014), which is predicted to have both nuclear localization and export signals, and later bound by the PETISCO/PICS complexes (Cordeiro Rodrigues et al., 2019; Zeng et al., 2019), which might be involved in 5′ end precursor processing, including decapping and 5′ end trimming. Subsequent 3′-end trimming is mediated by the exonuclease PARN-1 (Tang et al., 2016). The resulting mature *C. elegans* piRNAs are 21 nt small RNAs with a 5′ uracil bias (hence called 21U-RNAs). They also possess a 5′ monophosphate and a 3′ hydroxyl group at their extremities, and are post-transcriptionally 2′-O methylated at the 3′ end by the HENN-1 enzyme (Kamminga et al., 2010; Montgomery et al., 2012).

Mature piRNAs bind to PIWI subfamily proteins of the AGO family and guide them to complementary target transcripts. Two PIWI proteins with germline-restricted expression, PRG-1 and PRG-2, have been identified in *C. elegans*, although only PRG-1 is required for maintaining wild-type piRNA populations (Batista et al., 2008; Das et al., 2008; Wang and Reinke, 2008). These proteins localize to P-granules, specialized germline-specific phase-separated perinuclear structures that are the main site of piRNA-mediated post-transcriptional gene silencing activities (Updike and Strome, 2010).

Unlike in other animals, where PIWI/piRNA complexes can silence their targets via a PIWI endonuclease “slicing” activity, PRG-1/piRNA complexes in *C. elegans* silence their targets by triggering their degradation indirectly. The imperfect base pairing of the piRNA with its complementary transcripts elicits a localized silencing amplification response which targeted recruitment of the RNA-dependent RNA Polymerases (RdRPs) RRF-1 and EGO-1 leads to the generation of a secondary class of small RNAs that are 22 nt, have a 5′ guanine bias (hence called 22G siRNAs) and a 5′ triphosphate. Also termed secondary endogenous siRNAs (endo-siRNAs), these abundant molecules in turn promote robust and sustained silencing upon loading onto worm-specific Ago proteins (WAGOs) (Ashe et al., 2012; Bagijn et al., 2012a; Batista et al., 2008; Gu et al., 2009; Lee et al., 2012; Mao et al., 2015; Shirayama et al., 2012). *Mutator* family proteins MUT-16, MUT-7 and MUT-2 (Zhang et al., 2011), and the P-granules factor DEPS-1 (Suen et al., 2019) are also involved in this mechanism by allowing efficient 22G siRNAs amplification.

Interestingly, this silencing response can be inherited transgenerationally and last for several generations, even in *prg-1* mutant animals (Ashe et al., 2012). Once established, this long-term silencing is independent of the initial piRNA trigger and instead relies on the germline nuclear RNAi pathway, including the AGO protein HRDE-1 (Ashe et al., 2012; Buckley et al., 2012), the nuclear RNAi defective proteins NRDE-1/2-4 (Ashe et al., 2012), the nuclear RNA helicase EMB-4 (Akay et al., 2017), and several chromatin-associated factors, such as HPL-2, SET-25 and SET-32 (Ashe et al., 2012; McMurchy et al., 2017). Together, the piRNA and the germline nuclear RNAi pathways establish and maintain transgenerational silencing via the inheritance of small RNA populations and the deposition of chromatin silencing marks in the germline (Ashe et al., 2012; Buckley et al., 2012; Lee et al., 2012).

Since PRG-1/piRNA complexes tolerate several mismatches when selecting for complementary transcripts, they can theoretically target any sequence (Bagijn et al., 2012a; Lee et al., 2012; Shen et al., 2018; Zhang et al., 2018), including endogenous genes that are required to be expressed in the germline. To prevent silencing of germline-specific genes, some nematodes have evolved a protection mechanism involving the CSR-1/22G siRNA licensing pathway (Frøkjær-Jensen et al., 2016; Seth et al., 2013; Shen et al., 2018; Zhang et al., 2018), in which endogenous secondary 22G siRNAs associate with the Ago CSR-1 and prevent target recognition by the piRNA pathway. Given the importance of piRNAs in the maintenance of genome stability, it is essential to understand how they are generated. To date, many details about piRNA transcription are still unclear.

RNA Pol II transcribes piRNAs (Gu et al., 2012); (Billi et al., 2013). RNA Pol II is a holoenzyme that synthesizes a range of coding and non-coding transcripts through three precisely defined and controlled steps: initiation, elongation and termination (Adelman and Lis, 2012; Zhou et al., 2012). In eukaryotes, the RNA Pol II complex is typically composed of 12 subunits (RPB1-12), of which 10 constitute the catalytic core, and two (RPB4 and RPB7 in yeast) are required for transcription initiation (Cramer, 2004). RPB1/AMA-1, the largest subunit, is essential for polymerase activity through its carboxy terminal domain (CTD) and, in combination with RPB9, forms the DNA-binding groove of the holoenzyme (Acker et al., 1997). RPB2, the second largest subunit, constitutes a cleft in which the DNA template and the nascent RNA transcript are kept in proximity (Acker et al., 1992; Ponicsan et al., 2013), while RPB6 is part of a structure that stabilizes the active enzyme on the DNA template (Acker et al., 1994; del Río-Portilla et al., 1999; Wani et al., 2014). The structural role of the other subunits is less well documented.

RNA Pol II initiates transcription by assembling with general transcription factors to form the pre-initiation complex (PIC) at promoters. Once RNA synthesis commences, the PIC dissociates, enabling the polymerase to associate with elongation factors and form the elongation complex. This complex then “slides” on the DNA template, generating a complementary RNA molecule as it proceeds through the gene body. Faithful transcription is ensured by a proofreading mechanism called backtracking, during which the polymerase moves backwards on the DNA template to correct base mis-incorporation events (Bondarenko et al., 2006; Churchman and Weissman, 2011; James et al., 2017). As a result, the 3′ end of the nascent transcript is displaced from the enzyme′s active site, and the polymerase becomes transcriptionally inactive (Cheung and Cramer, 2011; Kettenberger et al., 2003; Komissarova and Kashlev, 1997; Nudler et al., 1997; Thomas et al., 1998; Wang et al., 2009). Backtrack recovery is required in order for the enzyme to resume transcriptional elongation, and often depends on cleavage of the backtracked RNA to generate a new 3′ end, which allows realignment with the active site (Chedin et al., 1998); (Kuhn et al., 2007); (Walmacq et al., 2009). Eukaryotic RNA Pol II has a weak intrinsic cleavage activity, which is strongly enhanced by the transcription elongation factor TFIIS ((Izban and Luse, 1992), and others).

RNA Pol II backtracking is associated not only with promoter-proximal pausing rescue and rapid transcription elongation, but also with transcription termination (Sheridan et al., 2019). Transcription termination is critical to determine borders between genes and to avoid interference with downstream-positioned loci, and it may be even more important for an organism like *C. elegans*, with a compact genome (C. elegans Sequencing Consortium, 1998). At non polyadenylated-type loci, such as those encoding for small nuclear RNAs (snRNAs), termination depends on specialized processing of the 3′ ends and often requires the Integrator complex, whose nuclease activity at specific cleavage sites is necessary for precursor transcript maturation (Baillat et al., 2005; Ezzeddine et al., 2011, 2012; Uguen and Murphy, 2003). Interestingly, transcription at motif-dependent piRNA loci shares evolutionary similarities to snRNA transcription (Beltran et al., 2019).

Here, by using a combination of genetics and biochemical approaches, we show that the RNA Pol II subunit RPB-9 is required to promote the Integrator-dependent cleavage of 3′ ends of nascent transcripts upon RNA Pol II backtracking for transcription termination at motif-dependent piRNA loci in *C. elegans*. This defect in transcription termination leads to a reduction in mature piRNA levels, which in turn results in a drastic depletion of HRDE-1-associated secondary 22G siRNAs and, ultimately, in the desilencing of two families of DNA transposons and a subset of somatic genes, likely in the germline.

## RESULTS

### *rpb-9* is required for piRNA-pathway integrity

In order to identify components of the piRNA pathway, we have described previously a forward genetics screen for animals defective in their ability to silence the “piRNA sensor”, a germline-specific transgene containing a *gfp* reporter and responsive to the endogenous piRNA 21UR-1 (Ashe et al., 2012; Bagijn et al., 2012a; Weick et al., 2014) (Figure 1A). Wild-type animals efficiently repress this *gfp* transgene via the piRNA pathway, while mutants of piRNA pathway components, such as *prg-1* and *mutator* class genes, fail to do so, and strongly desilence the piRNA sensor (Phillips et al., 2012). One of 22 independent mutants from our screen defined a new allele (*mj261*) of the gene *rpb-9*, which encodes for a conserved subunit of the DNA-dependent RNA Pol II enzyme (Cramer et al., 2000). Studies in yeast have proposed a role for Rpb9 in various steps of transcription, including start site selection and transcriptional elongation, and in transcription-coupled repair (Furter-Graves et al., 1994; Hemming et al., 2000; Li and Smerdon, 2002; Nesser et al., 2006). We decided to explore the possibility that *rpb-9* could provide a direct link between transcription and silencing in the context of piRNA-mediated repression in *C. elegans. C. elegans* RPB-9 is encoded on chromosome V and is composed of 167 amino acids. It comprises a central RPOL9 domain (a zinc ribbon) and a C-terminal zinc-finger motif-containing domain also found in the C-terminal portion of the transcription elongation factor TFIIS, hence named “TFIIS C”. The identified point mutation introduces a premature amber stop codon at glutamine 140 (CAG to TAG mutation), within the core of the TFIIS C domain (Figure 1B). As expected, *rpb-9* mutants do not express full-length RPB-9 protein at levels detectable via western blotting, although we cannot exclude residual RNA produced by transcriptional read-through (Figure S1A).

**Figure 1.**
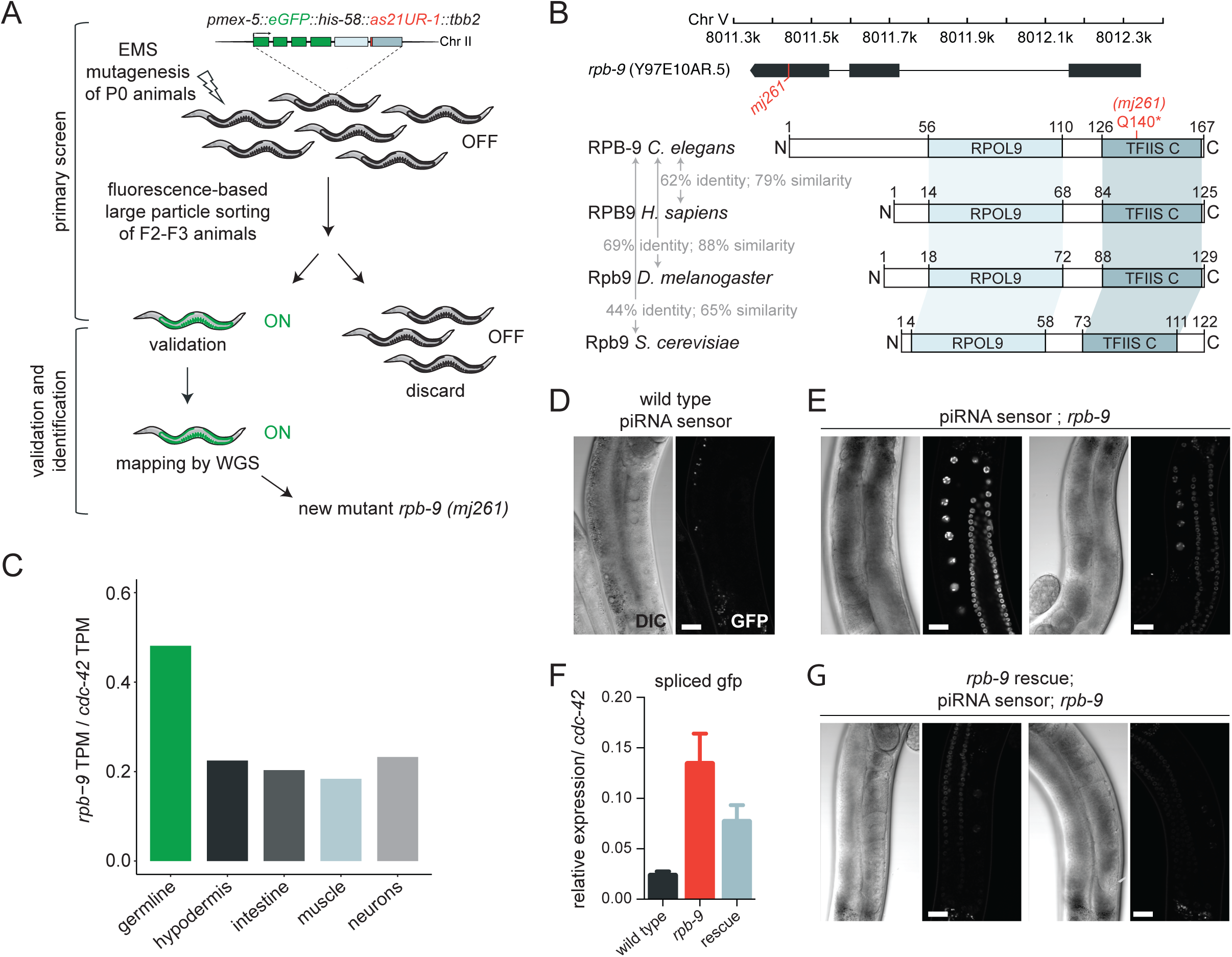
*rpb-9* is required for piRNA pathway integrity. **A**) Schematic of the EMS mutagenesis screen carried out using the piRNA sensor and identification of a novel allele *(mj261)* of the *rpb-9* gene. **B**) *rpb-9* gene locus, in scale (top) and comparisons of RPB-9 protein domains to homologs in other species (bottom). The introduced mutation is highlighted in red. **C**) Expression patterns of *rpb-9* in adult animals by RNA-sequencing. **D-G**) Quantification of piRNA sensor expression. Representative DIC and fluorescence microscopy images of piRNA sensor expression in wild type *(mjIs144)* (**D**), *rpb-9 (mj261)* (**E**) (two images with varying GFP expression are shown, scale bar = 20 µm) and validation of transgene expression by RT-qPCR (**F**). Representative DIC and fluorescence microscopy images of piRNA sensor expression in the *rpb-9* rescue *(mjSi70)* animals (two images with varying GFP expression are shown). Scale bar = 20 µm (**G**).

To explore the conservation of RPB9, we aligned the protein sequence of the *C. elegans* RPB-9 with its homologs in *Saccharomyces cerevisiae, Drosophila melanogaster* and *Homo sapiens*. We observed that both the RPOL9 and the TFIIS C domains are conserved in all these species (with sequence identities from 44% to 69%) and that *C. elegans* RPB-9 also possesses an additional N-terminal portion, which does not contain any known or conserved domains (Figure 1B and Figure S1B). We also compared the *C. elegans* RPB-9 and TFIIS sequences, and found that their respective C-terminal domains share 33% sequence identity and 53% sequence similarity, suggesting they might perform similar functions (Figure S1C). In *C. elegans, rpb-9* is expressed both in the soma and in the germline, consistent with its role as RNA Pol II subunit (Serizay et al., 2020) (Figure 1C). When visualized under a fluorescence microscope all *rpb-9* animals desilence the transgene in the germline, albeit at a variable level (Figure 1D, compare with Figure 1E). We measured the amounts of spliced *gfp* mRNA in these animals and observed increased levels of *gfp* expression, suggesting that *rpb-9* affects its targets prior to translation (Figure 1F).

To test whether the *rpb-9* mutation was causative for sensor desilencing, we constructed a synthetic transgene harboring the *rpb-9* coding sequence under control of a germline-specific promoter (*pmex5*::*rpb9::par-5, mjSi70*) and integrated it on chromosome I via the MosSCI technology (Frøkjaer-Jensen et al., 2008). This germline-specific allele restores wild-type levels of RPB-9 protein, as detected by western blotting (Figure S1A) and rescues the desilencing phenotype of *rpb-9* mutants (Figure 1F-G). Importantly, a similar transgene containing the Q140STOP mutation did not rescue the phenotype (Figure S1D). Altogether, these data demonstrate that *rpb-9* is required for the integrity of the piRNA pathway.

### *rpb-9* is required for establishing piRNA-mediated silencing

The piRNA pathway involves multiple steps and small RNA pathways, as well as an initiation and a transgenerational maintenance phase. In order to understand if *rpb-9* is required for the transgenerational maintenance of silencing, we crossed *rpb-9* animals with animals expressing a germline *gfp::h2b* reporter (Ashe et al., 2012; Buckley et al., 2012) and we assessed the transcriptional status of this transgene across generations upon a time-limited exposure to *gfp* RNAi (Figure 2A).

**Figure 2.**
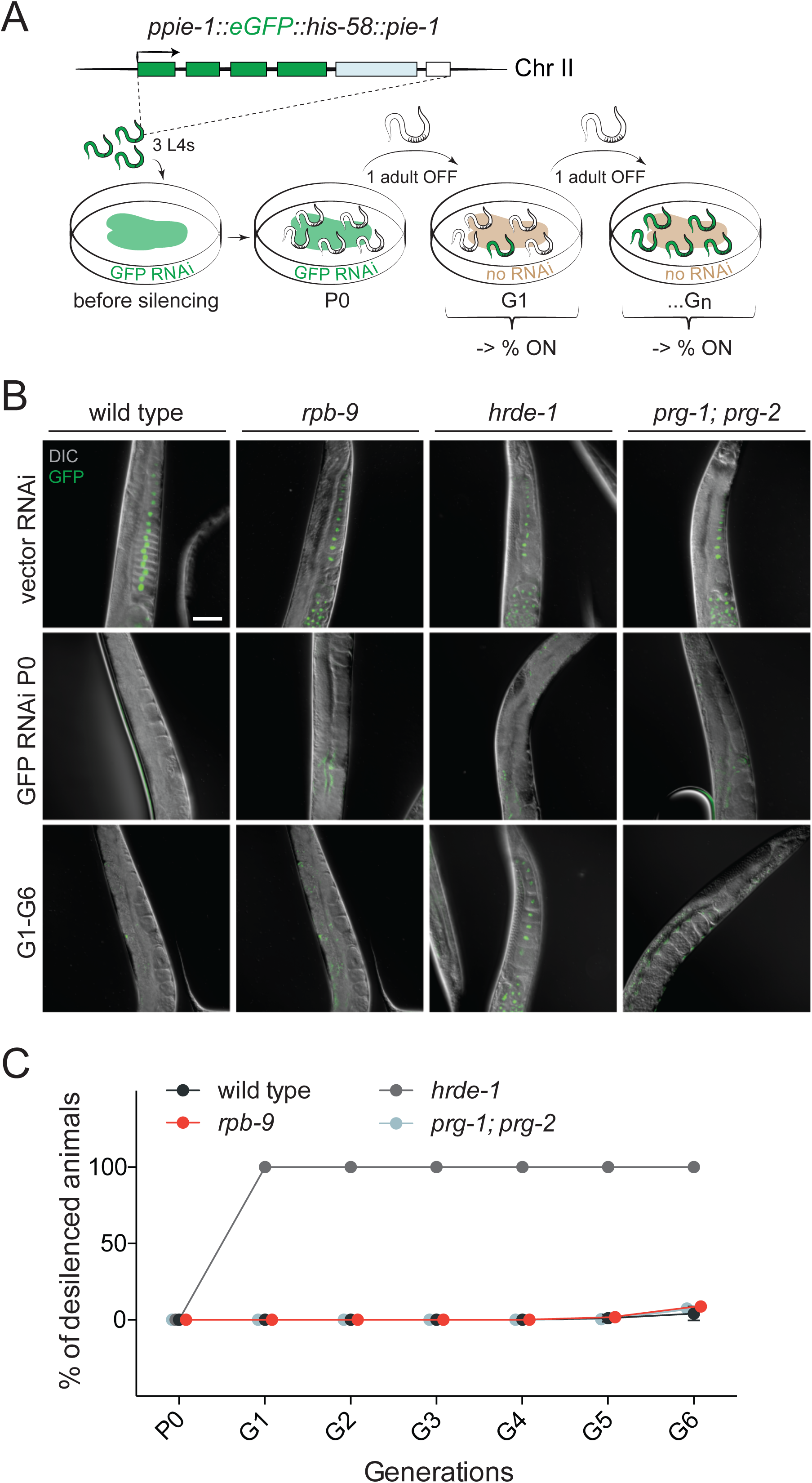
*rpb-9* is required for the initiation step of the piRNA pathway. **A**) Schematic of the germline-specific *pie1::gfp::h2b::pie-1* sensor (top) and schematic of the experiment (bottom). **B**) Representative fluorescence images of transgene expression in wild-type, *rpb-9 (mj261), hrde-1 (tm1200)* and *prg1 (n4357); prg-2 (n4358)* animals in the parental (P0) and inheriting (G1-G6) generations. Scale bar = 50 µm. **C**) Quantification of the percentage of desilenced individuals in wild-type, *rpb-9 (mj261)* and *prg-1 (n4357); prg-2 (n4358)* populations across generations.

In wild-type animals, exogenous double-stranded (ds) *gfp* RNA uptake for one generation is sufficient to establish and maintain reporter silencing for several generations (Ashe et al., 2012; Buckley et al., 2012). Conversely, mutants of components of the multi-generational germline nuclear RNAi pathway, albeit able to efficiently establish silencing in sustained presence of ds *gfp* RNA, fail to maintain it once this trigger is removed. This is the case for *hrde-1*, previously identified as the major germline-specific Ago that binds secondary endogenous 22G siRNAs and directs initiation of the transgenerational germline nuclear RNAi pathway (Figure 2B-C) (Ashe et al., 2012; Buckley et al., 2012). Like wild-type animals, *rpb-9* mutants can efficiently (100%) initiate silencing upon *gfp* RNAi, and are able to maintain it (91%-100%) even when the initial stimulus is removed (Figure 2B-C and Figure S2). This suggests that *rpb-9* is not required for exogenous RNAi and its inheritance. Instead it is required specifically for the endogenous piRNA mediated silencing. Similarly to *rpb-9* mutants, *prg-1; prg-2* double mutant animals are efficient in both the establishment and the maintenance of exogenous dsRNA-induced silencing (Figure 2B-C and Figure S2). Hence, we conclude that *rpb-9* is required for piRNA-dependent transgene silencing upstream of *hrde-1* and the multi-generational germline nuclear RNAi pathway, and independently of the exogenous-RNAi pathway.

### *rpb-9* is required to repress two DNA transposon families and a subset of somatic genes

In order to explore the genome-wide consequences of piRNA pathway disruption in *rpb-9* mutants, we performed a transcriptome analysis. To be able to capture all types of transcripts, including polyadenylated and non-polyadenylated RNAs, we prepared both total-RNA and poly-A-selected libraries from whole adult animals.

We first analyzed the expression of transposable elements. While no overall major changes were detectable in *rpb-9* mutants compared to wild type, we observed two transposon de-repression events, which affected two independent autonomous DNA transposon families, Chapaev-2 and CEMUDR1 (Figure 3A). Since DNA transposons are the most abundant transposable elements in *C. elegans*, and the only class suggested to be active in this organism (Bessereau, 2006; Laricchia et al., 2017), it was reasonable to assume that they would be bona-fide targets of piRNA-mediated silencing. When we scanned these two transposons families for potential piRNA target sites (Bagijn et al., 2012a) we found several matches. Chapaev-2 is a potential target of two piRNAs with up to two mismatches and ten piRNAs with up to three mismatches. CEMUDR1 is a potential target of one piRNA with perfect matching, five piRNAs with up to one mismatch and 17 piRNAs with up to three mismatches (Figure 3B). Moreover, we also checked for the density of endogenous 22G siRNAs antisense to these transposons in wild-type and *prg-1* animals and found that, for both transposons, 22G siRNAs were present at high levels in wild-type animals but were decreased in *prg-1* mutants. Together, these observations confirm that both Chapaev-2 and CEMUDR-1 are potential piRNA pathway targets and indicate that *rpb-9* is required to repress their activity in the germline.

**Figure 3.**
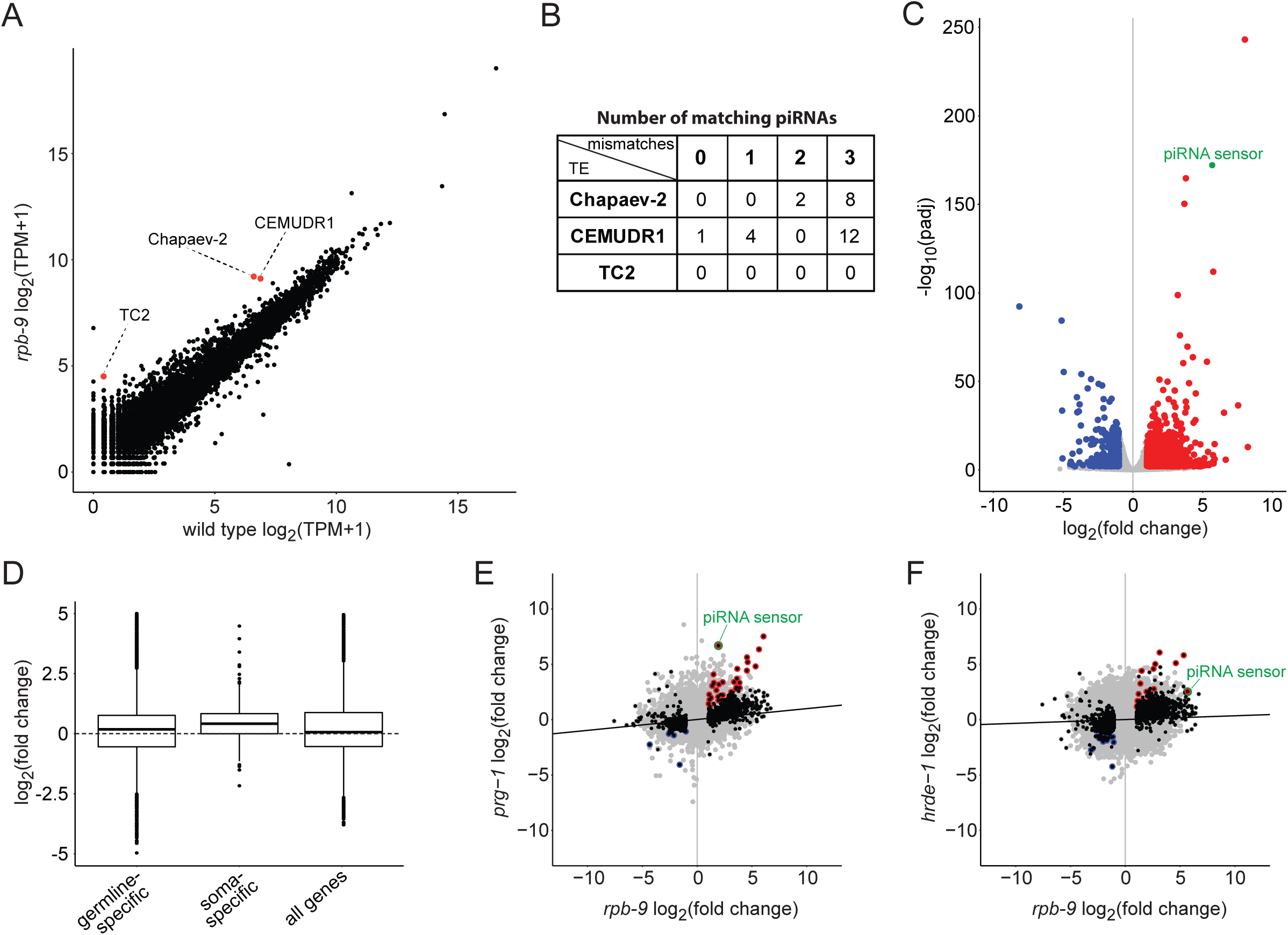
*rpb-9* represses two DNA transposon families and a subset of somatic genes. **A**) Differential expression analysis of transposable elements in *rpb-9* (*mj261*) mutants versus wild type (polyA-selected RNA-seq libraries). **B**) Table showing the numbers of matching piRNAs (with 0, 1, 2 and 3 mismatches respectively) for each of the upregulated transposon families. **C**) Genome-wide differential expression analysis of *rpb-9* (*mj261*) mutants versus wild-type (polyA-selected RNA-seq libraries). [p<=0.01, log_2_(fold change)>1 or <-1]. **D**) Differential expression analysis of germline-specific and soma-specific genes, as classified in (Reinke et al., 2004). “All genes” is shown for comparison. (PolyA-selected RNA-seq libraries). **E-F**) Pairwise correlation plots of *rpb-9 (mj261)* and *prg-1 (n4357)* (**E**) or *hrde-1 (tm1200)* (**F**) transcriptomes. Differentially expressed genes in *rpb-9 (mj261)* are shown in black, shared upregulated genes in red, shared downregulated genes in blue. The piRNA sensor transcript is highlighted in green. (Total RNA “Ribo-Zero” RNA-seq libraries).

In *C. elegans* the piRNA pathway has been previously implicated in the regulation of endogenous genes (Rojas-Ríos and Simonelig, 2018). For this reason, we also analyzed the expression of endogenous mRNA transcripts in *rpb-9* mutants. When we compared the polyA-selected transcriptome of *rpb-9* and wild-type animals, we found a total of 1,556 misregulated transcripts in *rpb-9* mutants, with 292 being downregulated (18.76%) and 1,264 upregulated (81.23%) (Figure 3C).

Since the piRNA pathway acts in the germline, we wondered whether some of the upregulated transcripts in *rpb-9* mutants corresponded to desilenced somatic genes in the germline as is the. To test this, we took advantage of published data (Bezler et al., 2019); (Reinke et al., 2004) to classify transcripts into “germline-specific” and “soma-specific” and used these classes to filter deregulated genes in *rpb-9* mutants. We found that, while germline-specific transcripts did not vary significantly, soma-specific transcripts tended to be upregulated in *rpb-9* mutants (Figure 3D). This suggests that, other than in controlling transposon activity, *rpb-9* might be involved in the repression of a subclass of somatic genes in the germline. Importantly, analysis of total-RNA libraries yielded comparable results, both for transposable elements and mRNA transcripts (Figure S3A-C).

In order to explore the relationship between *rpb-9* and the other major components of the piRNA pathway in the control of germline integrity, we compared the transcriptomes of our *rpb-9* mutants with those of *prg-1* and *hrde-1* mutants. We observed that *rpb-9* and *prg-1* mutants share a total of 51 deregulated genes (22 upregulated and 29 downregulated) (Figure 3E), while *rpb-9* and *hrde-1* share 42 (36 upregulated and 6 downregulated) (Figure 3F). Importantly, we found the piRNA sensor transcript as a commonly upregulated target in both *rpb-9* vs *prg-1* and *rpb-9* vs *hrde-1* comparisons, consistent with the idea that *rpb-9* represses germline transgenes via both the piRNA pathway and its downstream effector, the nuclear RNAi pathway. These observations suggest that *rpb-9* is required, together with *prg-1* and *hrde-1*, for gene silencing at a subset of piRNA pathway targets.

In order to refine this analysis we decided to focus specifically on germline transcriptomes. By taking advantage of published datasets from gonad-dissected samples (Reed et al., 2020) and comparing them with our own, we asked what proportion of misregulated genes in *rpb-9* mutants could originate from within the germline. We observed a significant overlap between upregulated genes in *rpb-9* animals and upregulated genes in *prg-1* mutant germlines (525 genes, 40.8 %, r^2^ > 0.5) (Figure S3D), in agreement with the hypothesis that transcriptome misregulations in *rpb-9* mutants mostly arise from germline-related piRNA pathway defects.

### Many upregulated genes show invariant or reduced RNA Pol II enrichment

We reasoned that the downregulated genes would likely be a result of a defect in the canonical role for *rpb-9* in promoting transcription as part of the RNA Pol II complex, consistent with ubiquitous *rpb-9* expression. Thus, in order to decipher the role of *rpb-9* in the context of the piRNA pathway, we decided to focus our attention on upregulated genes. Given the reported role for *rpb-9* in TSS selection (Ghazy et al., 2004; Hull et al., 1995; Sun et al., 1996), we sought to understand if upregulation/desilencing of these genes would merely depend on increased binding of RNA Pol II at their TSS, or if another mechanism could be implicated. In order to explore these possibilities, we performed an RPB-1/AMA-1 Chromatin Immunoprecipitation followed by sequencing (ChIP-seq) analysis. We did not observe major differences in RNA Pol II binding genome-wide (Figure S4A), but noticed differential binding patterns for upregulated genes in *rpb-9* mutants compared to wild type. A certain proportion (138 genes, 33.3%) of the 384 upregulated genes with detectable RNA Pol II signal also showed increased RNA Pol II binding at the TSS and within the gene body, compared to wild type (Class I genes) (Figure 4A). Surprisingly however, we observed that the majority of upregulated genes (246 genes, 66.7%) displayed unchanged or reduced RNA Pol II binding in *rpb9* animals, despite being upregulated (or desilenced) (Class II (Figure 4B) and Class III genes (Figure 4C)). Importantly, the piRNA sensor locus belonged to class III genes (Figure 4D). Analysis of total RNA libraries yielded similar results (Figure S4B-D). These results suggested an additional role for *rpb-9*, possibly uncoupled from its canonical function as RNA Pol II subunit at a subset of endogenous genes.

**Figure 4.**
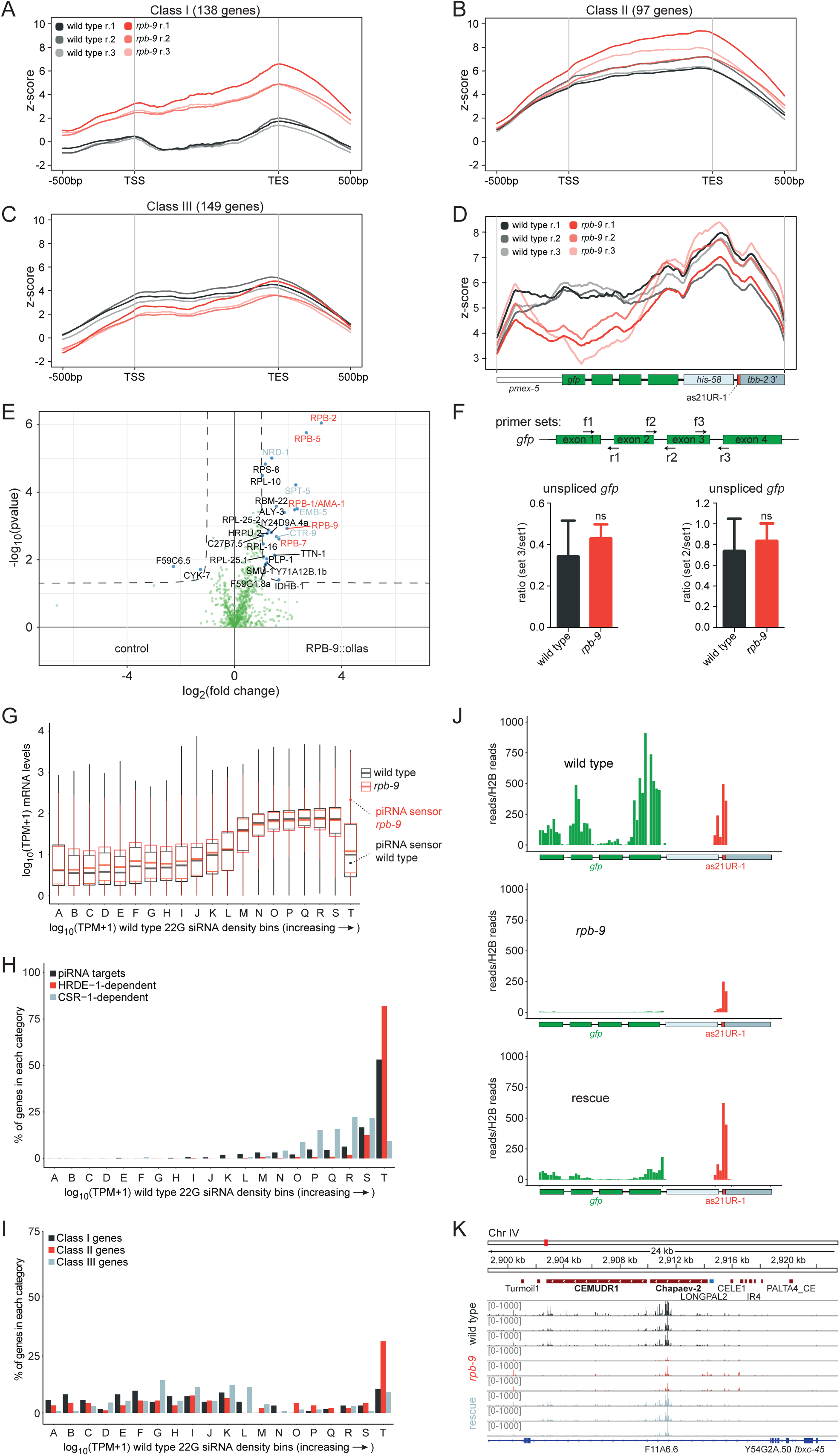
*rpb-9* is required for the robust production of piRNA-pathway dependent HRDE-1-associated secondary 22G siRNAs. **A-C**) Analysis of RNA Pol II binding (RPB-1/AMA-1 ChIP-seq) at upregulated genes in *rpb-9 (mj261)* mutants, as defined in Figure 3C. Class I: upregulated genes with increased RNA Pol II binding (**A**); Class II: upregulated genes with invariant RNA Pol II binding (**B**); Class III: upregulated genes with reduced RNA Pol II binding (**C**). n=3 biological replicates are shown per each genotype. (PolyA-selected RNA-seq libraries) **D**) RNA Pol II binding profile over the piRNA sensor locus. **E**) Enrichment analysis of RPB-9-bound proteins in *rpb-9::ollas (mj604)* versus wild type by IP/MS. [p<=0.05, log_2_(fold change)>1 or <-1]. n=4 technical replicates (pooled). **F**) Transgene transcript length is not affected in *rpb-9 (mj261)* mutants. Scheme of the *gfp* unspliced transcript showing the positions of RT-qPCR primers (top) and pairwise ratios (3′ to 5′) of unspliced *gfp* amounts along the transcript as indicated by the primer pairs (bottom). ns= not significant (two-tailed t-test). **G**) Transcriptome binning according to increasing 22G siRNA density in wild-type animals (grey). Mean normalized *rpb-9 (mj261)* mRNA reads (red) are overlaid with mean normalized wild-type mRNA reads (grey). (PolyA-selected RNA-seq libraries) The piRNA sensor transcript is indicated with red (*rpb-9 (mj261)*) and grey (wild type) dots. **H**) Distribution of piRNA targets (as defined in (Bagijn et al., 2012a) and of HDRE-1- and CSR-1-dependent 22G siRNAs across bins as defined in G). **I**) Distribution of Class I, Class II and Class II genes (as defined in Figure 4A-C) across bins as defined in G). **J**) Distribution of normalized 22G siRNA reads mapping over the piRNA sensor. **K**) Distribution of normalized reads mapping over the CEMUDR1 and Chapaev-2 DNA transposons.

In order to elucidate novel *rpb-9* nuclear functions, we set out to identify its protein interactors. To be able to perform an immunoprecipitation followed by mass spectrometry (IP-MS), we generated an endogenously-tagged *rpb-9::ollas* strain using the CRISPR/Cas9 system (Akay et al., 2017; Jiang and Marraffini, 2015). RPB-9::OLLAS is expressed (Figure S4E) and does not induce piRNA sensor desilencing (Figure S4F), suggesting that the protein fusion is functional.

In the pull-down experiment, we found a total of 25 proteins with significantly enriched binding compared to the control (Figure 4E, Welch t-test, p<0.05). Four of them were other polymerase subunits: RPB-1/AMA-1, already reported to be associating with RPB-9 to form the DNA-binding groove of the holoenzyme in human (Acker et al., 1997), and RPB-2, RPB-5 and RPB-7, all contacts that were never reported before. Interestingly, other proteins were factors involved in transcription elongation and termination, including the elongation factors SPT-5 (a component of the DSIF complex), CTR-9 (a component of the PAF-1 complex) and EMB-5 (human SUPT6h homolog), and the early-termination factor NRD-1 (human SCAF4 homolog). The identity of these binding partners is consistent with a previously described role for Rpb9 in transcriptional elongation in yeast (Awrey et al., 1997; Hemming et al., 2000; Van Mullem et al., 2002).

Given these results, we hypothesized that *rpb-9* could be important for transcriptional elongation of class II and class III genes, and we speculated that premature termination of transcriptional elongation could manifest in *rpb-9* mutants, result in shorter nascent transcripts and hence in a limited docking platform for the HRDE-1/22G-siRNA silencing machinery. In order to test this hypothesis, we designed primers spanning three intron-exon junctions along the *gfp* transgene within the piRNA sensor and measured unspliced *gfp* levels in *rpb-9* mutants and wild-type animals by real-time qPCR. The results show that, although the unspliced transcript is significantly upregulated in *rpb-9* mutants, the ratios of unspliced *gfp* levels along the transcript (3′ end vs 5′ end) are not significantly different compared to wild type, suggesting that transcriptional elongation at the piRNA sensor locus is in fact efficient in *rpb-9* animals (Figure 4F). Taken together, these results suggest that piRNA-dependent silencing establishment in *rpb-9* mutants must be defective upstream of nascent transcript synthesis.

### *rpb-9* is required for robust secondary piRNA pathway-dependent HRDE-1-associated 22G siRNA production

We tested whether the defect in piRNA target silencing in *rpb-9* mutants was caused by a decrease in the abundance of endogenous secondary 22G siRNAs. Indeed, a strong reduction in the amount of these molecules could explain why reduced or unchanged RNA Pol II binding on class II and III genes (with wild-type lengths of nascent transcripts) still results in their upregulation/desilencing.

Endogenous secondary 22G siRNAs are not only mediators of silencing, but they can also counteract it. This is for example the case for CSR-1-associated 22G siRNAs, which are the effectors of a protection mechanism against the piRNA-mediated silencing of germline-expressed genes (Wedeles et al., 2013). We reasoned that, for these types of targets, high 22G siRNA amounts would correlate with higher gene expression rates. On the other hand, piRNA target genes would display high 22G siRNA amounts but low expression rates. For these reasons, we explored the correlation between mRNA expression levels and endogenous secondary 22G siRNA abundance genome-wide. We subdivided the transcriptome into 20 bins (A to T), each containing 857 genes, which were ranked according to their 22G siRNA density in wild type (Figure 4G). We observed a general trend where 22G siRNA density correlated with mRNA expression levels, consistent with the idea that the more an mRNA is expressed, the more 22G siRNAs can be produced for that locus. A notable exception was the last bin (T), corresponding to the genes with the highest 22G siRNAs density (top 5%), but which displayed lower mRNA expression levels. This pattern is consistent with active 22G siRNA-mediated silencing at high 22G siRNA densities. Interestingly, the mean expression levels of genes in bin T was higher in *rpb-9* mutants compared to wild type. This suggests that these genes are likely direct piRNA targets. In further support of this hypothesis, we found that the piRNA sensor belongs to this subset of genes (bin T). Finally, we observed that the highest proportion of piRNA targets (with ≤1 mismatch, (Bagijn et al., 2012a)) indeed belong to this bin (Figure 4G and Figure 4F).

Since different Ago proteins bind to different subsets of 22G siRNAs and mediate different outcomes with regard to gene silencing and gene licensing, we then examined how HRDE-1-and CSR-1-dependent 22G siRNAs were distributed across the bins (Figure 4H). We observed that, where 22G siRNAs abundance and mRNA expression correlated the most (bins N to S), CSR-1-associated 22G siRNAs were present at the highest proportions and the mean expression of the corresponding genes did not change significantly in *rpb-9* mutants (two sample t-test, p<0.05). Conversely, where 22G siRNAs abundance and mRNA expression were anticorrelated (high 22G siRNAs but low mRNA expression) (bin T), HRDE-1-associated 22G siRNAs were found in the highest proportions and the mean expression of the corresponding genes was higher in *rpb-9* mutants compared to wild type, once again suggesting that the genes belonging to this class could be real *rpb-9*-dependent piRNA targets. Interestingly, when we analyzed how class I, class II and class III genes were partitioned across the bins, we observed that class II (and to a minor extent class III) genes mostly resided in the last bin (T), in agreement with our hypothesis of them being direct downstream targets of *rpb-9*-dependent piRNAs (Figure 4I). Importantly, these observations held true also after analysis of total-RNA libraries (Figure S4G-I).

As the piRNA sensor belonged to bin T, we examined the distribution of antisense 22G siRNAs mapping to the *gfp* transgene more closely. We observed that, while *gfp* 22G siRNAs were present at high levels over all exons in wild-type animals, they were mostly depleted in *rpb-9* mutants and partially recovered in the rescue strain (Figure 4J), indicating that *rpb-9* is indeed required to induce the generation of secondary 22G siRNAs targeting the piRNA sensor in amounts sufficient to repress it. We observed a similar trend in secondary 22G siRNAs mapping over the two desilenced transposons Chapaev-2 and CEMUDR1, confirming that *rpb-9* is required to suppress their expression via the piRNA-dependent generation of 22G siRNAs (Figure 4K). Next, we quantified the density of 22G siRNAs mapping over all piRNA targets (Bagijn et al., 2012a), and observed that a significant proportion of these transcripts (33%, 64 genes) were depleted in antisense 22G siRNAs in *rpb-9* mutants compared to wild type (hypergeometric test, p<0.017) (Figure S4J). This confirms that *rpb-9* is required for the production of 22G siRNAs at a subset of piRNA targets. We observed a similar phenotype in *hrde-1* mutants, although the affected targets did not overlap completely with those of *rpb-9* animals, in agreement with our transcriptome correlation analysis (Figure 3F). Consistent with a prominent role for *prg-1* in the generation of piRNAs, *prg-1* mutants showed an even stronger depletion in 22G siRNAs at piRNA targets. Importantly, the defect of *rpb-9* mutants was reverted in the *rpb-9* rescue strain (Figure S4J). Together, these data indicate that *rpb-9* is required to produce levels of HRDE-1-bound 22G siRNAs that are sufficient to efficiently establish silencing at piRNA target loci.

### *rpb-9* is required for the generation of mature piRNAs

In wild-type animals, high levels of endogenous 22G siRNAs are part of a self-sustaining response initiated by RdRPs upon binding of primary siRNAs, including piRNAs, to complementary transcripts. We therefore decided to measure the amounts of mature piRNAs in *rpb-9* mutants.

First, we analyzed global mature piRNA levels and found a significant reduction of their expression in *rpb-9* mutants, which was restored in the rescue strain (Figure 5A, two sample t-test p<0.05). Although this reduction was not as strong as in *prg-1* mutants (Suen et al., 2019), it suggests that *rpb-9* is required to produce mature piRNAs. We next wondered whether this reduction in expression levels affected all piRNAs or only a subset of them. We thus plotted piRNA expression data along chromosome coordinates in equal bins, and observed that all the piRNA species encoded in the two chromosome IV clusters were downregulated in *rpb-9* mutants compared to wild type (Figure 5B) but recovered in the rescue (Figure S5A). This suggests that *rpb-9* controls piRNA production via a single mechanism, possibly independent of sequence variations among single piRNAs.

**Figure 5.**
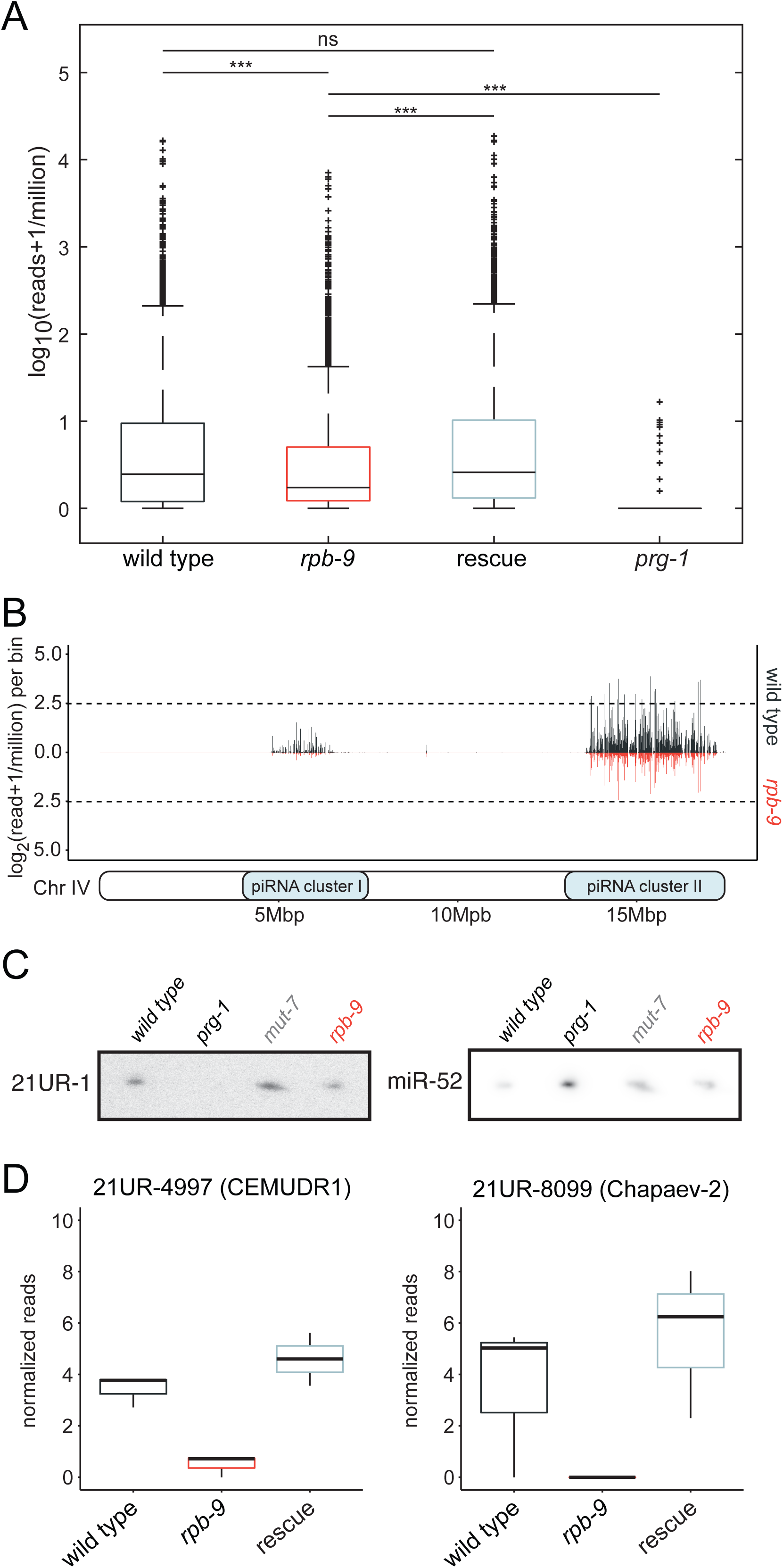
*rpb-9* is required for the generation of mature piRNAs. **A**) expression levels of mature piRNAs in wild-type, *rpb-9 (mj261), rpb-9* rescue and *prg-1 (n4357)* animals. **B**) Mature piRNA expression along chromosome IV coordinates (motif-dependent piRNA clusters I and II) in wild-type and *rpb-9 (mj261)* animals. **C**) Northern blot quantification of mature piRNA 21UR-1 levels in wild-type, *prg-1 (n4357), mut-7 (mj253)* and *rpb-9 (mj261)* animals. **D**) Quantification of the levels of the mature piRNAs 21UR-4997 and 21UR-8099 by small RNA sequencing (raw reads normalized to library size).

We also quantified the abundance of piRNA 21UR-1, whose complementary sequence is present within the piRNA sensor. We observed a subtle reduction in the mature levels of this piRNA in *rpb-9* mutants compared to wild type, both by Northern blotting (Figure 5C) and RT-qPCR (Figure S5B). Although this decrease is not statistically significant, it nevertheless is likely the primary cause of the observed loss of 22G siRNAs. In agreement with this hypothesis, we observed that, although some 22G siRNAs were still detectable over the 21UR-1 target site within the piRNA sensor (Figure 4J), their levels were lower than wild type, and 22G siRNA spreading to the rest of the transcript was defective. We observed this previously in other piRNA pathway mutants (Akay et al., 2017).

Similarly, the amounts of some of the piRNAs predicted to target the Chapaev-2 and CEMUDR-1 transposons were decreased in *rpb-9* mutants (Figure 5D), confirming that *rpb-9* represses them via the piRNA pathway. All together, these data indicate that *rpb-9* is required to generate wild-type levels of mature piRNAs.

### *rpb-9* is required for transcription elongation/termination at motif-dependent piRNA loci

We reasoned that this decrease in the abundance of mature piRNAs in *rpb-9* mutants would originate from a defect in the production of the corresponding piRNA precursors, so we decided to investigate the role of *rpb-9* in piRNA loci transcription. To do so, we sequenced short capped RNAs from isolated germ nuclei obtained from young adult animals, after separation of chromatin and nucleoplasmic fractions (as described in Beltran et. al., 2020, co-submitted). In *rpb-9* mutants, capped piRNA precursors were longer when compared to wild-type and rescue controls (Figure 6A). Similarly, the length distribution of nascent piRNA precursors, which follows a bimodal distribution (Beltran et al., 2020, co-submitted), was shifted also in *rpb-9* mutants for both peaks of nascent RNA lengths (Figure 6B-C). This peak-shift phenotype is identical to the one observed in *tfiis* mutants (Beltran et. al., 2020, co-submitted).

**Figure 6.**
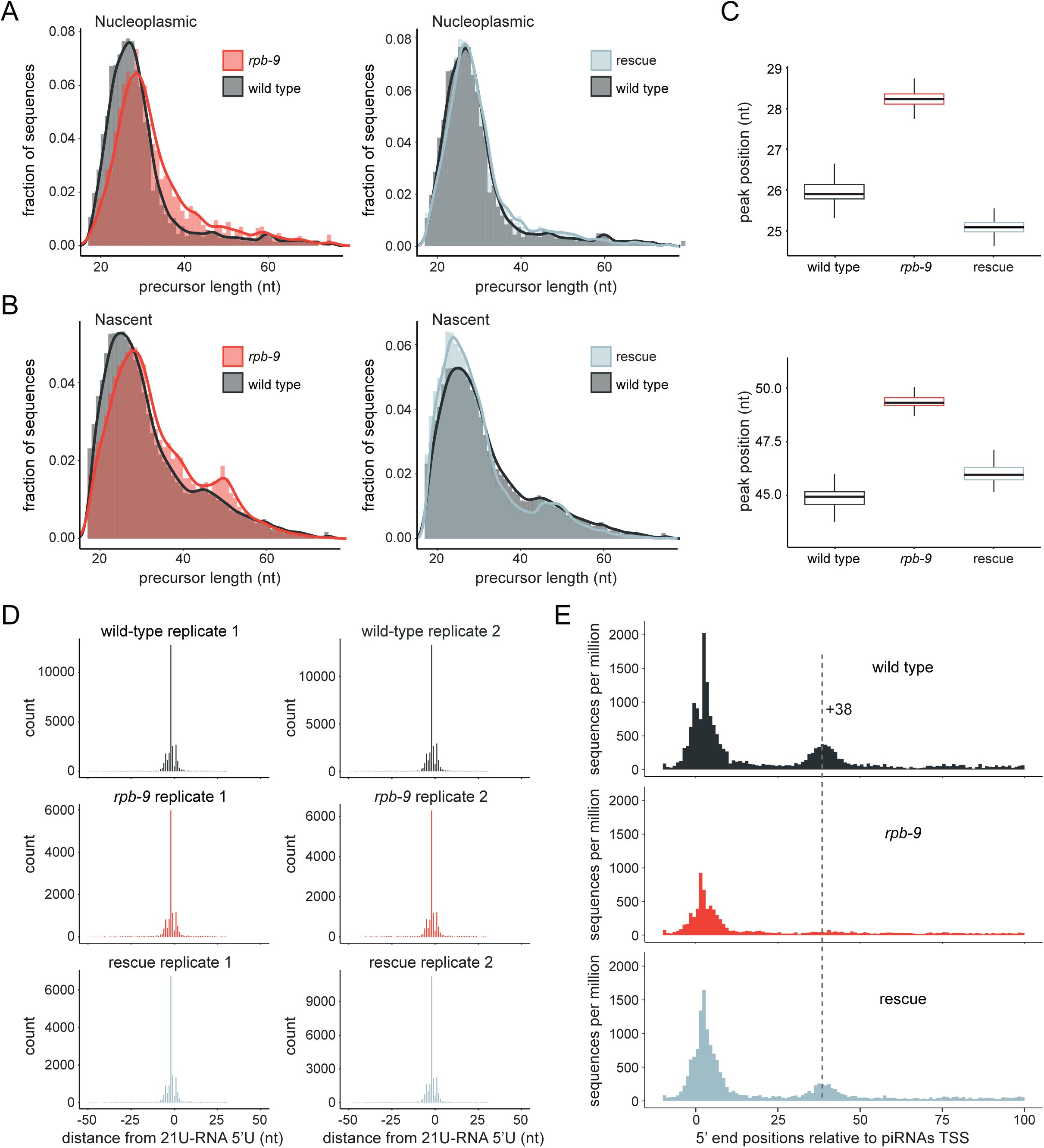
*rpb-9* is required for transcription elongation/termination at piRNA loci. **A**) Distributions of nucleoplasmic piRNA precursor lengths in *rpb-9 (mj261)* and rescue *(mjSi70)* animals compared to wild type. **B**) Distribution of nascent piRNA precursor lengths in *rpb-9 (mj261)* and rescue *(mjSi70)* animals compared to wild type. **C**) Peak positions of nascent piRNA precursor length distributions in 2000 subsamples of 5000 precursor sequences sampled without replacement. **D**) Distribution of 5′ ends of short capped RNA reads mapping to motif-dependent piRNA loci. Total counts of read 5′ ends mapping within a 50bp window of motif-dependent piRNA loci, aggregated by position relative to 21U-RNA 5′ U sites. A comparable enrichment of reads initiating 2 nt upstream of 21U-RNA 5′ U sites is observed across replicates and genotypes. **E**) Positions of unique 5′ monophosphate small RNA reads mapping at piRNA promoters after removal of reads >15 nt initiating at annotated piRNA 5′ U sites. The average signal is normalized to sequences per million of mapped reads.

Previous studies in yeast have shown that Rpb9 both modulates the selection of the TSS and is involved in the elongation of transcription: in cells lacking Rpb9, the population of TSSs is shifted upstream at a subset of promoters (Furter-Graves et al., 1994; Hull et al., 1995; Sun et al., 1996) and, in *in vitro* elongation assays, an RNA Pol II complex lacking Rpb9 pauses at intrinsic elongation blocks at a lower frequency compared to wild type (Hemming et al., 2000). In order to test whether TSS selection at piRNA loci depends on RPB-9, we analyzed the distribution of the 5′ ends of short capped RNA reads. For both wild-type and *rpb-9* animals, as well as for rescue individuals, we observed an enrichment of reads initiating 2 nt upstream of piRNAs U sites, suggesting that TSS selection is not impaired in *rpb-9* mutants and that the increase in piRNA precursor length is rather due to a defect in elongation/termination at the 3′ end (Figure 6D). This role is consistent with the IP-MS data (Figure 4E), which indicate that RPB-9 strongly interacts with components of the elongation machinery.

Together, these data indicate that RPB-9 is required for cleavage of the 3’ ends of nascent RNAs upon RNA Pol II backtracking, which is stimulated by TFIIS, promoting the resumption of transcription elongation. At piRNA loci, termination of promoter-proximal RNA Pol II depends on the 3′ end cleavage activity of the Integrator complex (Beltran et al., 2020, co-submitted). In order to understand if *rpb-9* could be required to recruit Integrator activity, we examined the presence of ∼20 nt long 3′ nascent RNA cleavage fragments downstream of 21U-RNAs in our samples. Interestingly, we observed a marked decrease in the abundance of these fragments in *rpb-9* mutants (Figure 6E and Figure S6), as found in a *tfiis* mutant background (Beltran et. al., 2020, co-submitted). Altogether, these data suggest that RPB-9 is important for Integrator-mediated termination of promoter-proximal RNA Pol II at motif-dependent piRNA loci.

## DISCUSSION

Initiation of piRNA pathway-mediated repression depends on the faithful transcription of piRNA loci. Here, we have characterized the contribution of the RNA Pol II subunit RPB-9 to this process. We have shown that *rpb-9* is required to promote efficient transcription elongation/termination at motif-dependent piRNA loci by eliciting Integrator-dependent cleavage of the 3′ end of nascent transcripts, and demonstrated that this activity is required to exert proper repression of two DNA transposon families, as well as a subset of somatic genes, in the germline. Mechanistically, a functional RPB-9 protein is required to produce a sufficient amount of mature piRNAs to ensure the generation of high levels of HRDE-1-dependent secondary 22G siRNAs, which target complementary transcripts and thereby induce their silencing (Figure 7).

**Figure 7.**
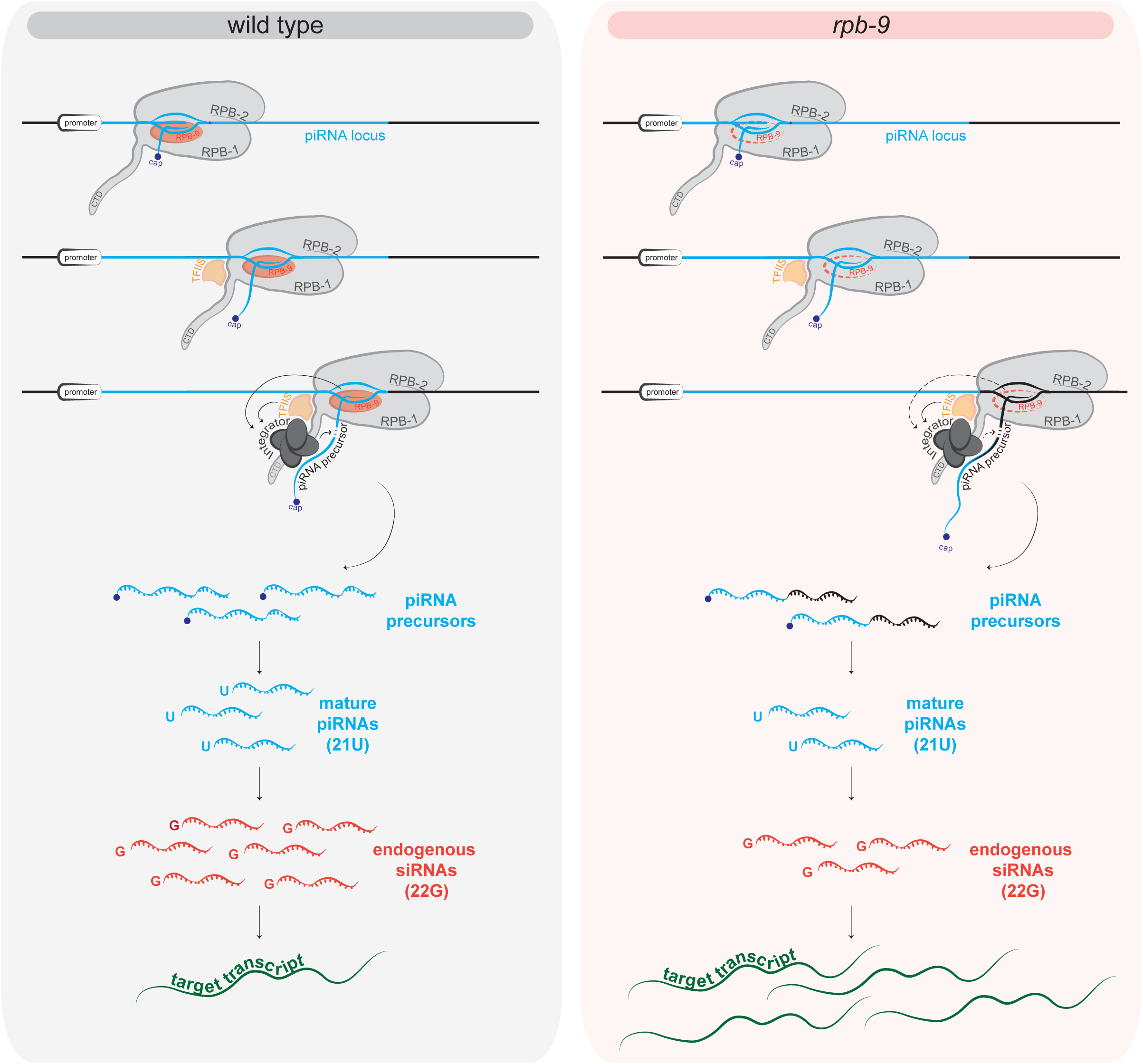
Model of RPB-9 function. In wild-type animals, RPB-9 promotes Integrator-dependent cleavage of nascent piRNA precursors. piRNA precursors are then processed into mature piRNAs, which trigger a potent 22G siRNAs amplification response. As a result, complementary target transcripts are silenced. In *rpb-9* mutants, transcription termination is defective and leads to longer piRNA precursors. As a consequence, the amount of mature piRNAs is decreased and the amplification response impaired. Target transcripts fail to be silenced.

Interestingly, a relatively modest decrease in mature piRNA levels in *rpb-9* mutants is sufficient to provoke a drastic reduction in the amounts of endogenous secondary 22G siRNAs antisense to piRNA target loci. This is consistent with the fact that the piRNA pathway relies heavily on amplification mechanisms and small RNA thresholds to establish silencing. Because of this, it is possible that genes targeted by low abundance piRNAs are more sensitive to small changes in their amounts, and hence more susceptible to desilencing. Indeed, this seems to be the case for the Chapaev-2 and CEMUDR1 DNA transposon families, which are targeted by lowly-abundant piRNAs and show a strong desilencing phenotype in our mutants. Chapaev-2 and CEMUDR1 are also desilenced in *prg-1* (Bagijn et al., 2012a) *and hrde-1* (Akay et al., 2017) *mutants, in agreement with our conclusion that rpb-9* functions upstream of both these components in the piRNA pathway. However, this is not the case in *prde-1* mutants; *prde-1*, like *rpb-9*, is also required for the transcription of motif-dependent piRNA loci (Weick et al., 2014). We speculate that this discrepancy is due to the fact that PRDE-1 and RPB-9 perform different molecular functions in the context of piRNA loci transcription. While PRDE-1 defines the site of piRNA precursor generation, thereby influencing transcription initiation, RPB-9 is rather required for the elongation and termination phases. As a consequence, it is possible that PRDE-1 and RPB-9 control different subsets of piRNA transcription units and/or impact the same loci to different extents. As mentioned above, piRNA-dependent transposon repression relies on specific small RNA thresholds: in this particular case, even minor differences in the abundance of certain mature piRNAs in *rpb-9* vs *prde-1* mutants could result in a profound effect on the repression of the same target.

An interesting observation is that the elongation/termination defects of *rpb-9* mutants are not detectable at deregulated genes other than at piRNA loci, suggesting that RPB-9, a conserved RNA Pol II subunit, may fulfill a very specific role in *C. elegans*. Perhaps RPB-9 is part of an ancient network which, together with SNPC-4 (Kasper et al., 2014) and the USTC complex (Weng et al., 2019), has been recruited to direct transcription at piRNA loci. The recent discovery that motif-dependent piRNA units share evolutionary similarities with the highly conserved snRNA genes (Beltran et al., 2019), together with the fact that transcription of both these types of loci depends on the Integrator complex (Baillat et al., 2005; Ezzeddine et al., 2011, 2012; Uguen and Murphy, 2003) (Beltran et al., 2020 co-submitted), seem to favor this hypothesis. Whether *rpb-9* is required for transcription of snRNA genes in *C. elegans* is however currently unknown.

Studies in yeast have shown that Rpb9 is located at the tip of the “jaws” of the RNA Pol II complex, proposed to clamp the DNA downstream of the enzyme active site (Cramer et al., 2001; Gnatt et al., 2001). Despite this remarkable feature, deletion of Rpb9 in yeast does not lead to cell death (Woychik et al., 1991), but only to minor transcriptomic changes in metabolism-related genes (Hemming et al., 2000), suggesting that Rpb9 is an accessory subunit in this organism. To understand if this was the case also in *C. elegans*, we initially set out to generate complete knockout mutant animals via the CRISPR/Cas9 technology. Although we were able to observe heterozygous editing events upon sgRNA injections in the germline, our attempts at recovering *rpb-9* homozygous knockout mutants in the F2 progeny failed (data not shown). This suggests that, similarly to rpb9 in *D. melanogaster, rpb-9* could in fact be essential in *C. elegans*. Although further experiments are required to confirm this hypothesis, we speculate that the role of RPB-9 in the piRNA pathway, which is absent in yeast, is what makes this protein so important in *C. elegans*.

Here we have shown that *S. cerevisiae* Rpb9 and *C. elegans* RPB-9 share sequence identity at the two identified domains. In yeast, the N-terminal domain of Rpb9 has been shown to influence TSS selection, while the C-terminal domain has been reported to be involved in intrinsic transcript cleavage (Hemming et al., 2000). In our *rpb-9* mutants, a truncated but partially functional RPB-9 protein might explain why TSS selection is not affected, but transcriptional elongation/termination is. Indeed, our mutation is supposed to interrupt translation within the TFIIS C domain, but leaves the RPOL9 domain intact, suggesting that the both RPOL9 and TFIIS C domains may perform conserved functions in *S. cerevisiae* and *C. elegans*.

The conservation between the C-terminal domains of Rpb9 and tfiis in yeast has prompted *in vitro* studies with purified proteins. These early reports show that RNA Pol II ternary complexes lacking Rpb9 are defective in their response to TFIIS-stimulated readthrough past an elongation block. They also demonstrate that the absence of Rpb9 does not affect the intrinsic cleavage ability of the RNA Pol II nor its binding to TFIIS, and that the addition of purified Rpb9 *in trans* restores the response of the complex to TFIIS (Awrey et al., 1997). It was thus proposed that Rpb9 might transmit a molecular signal from TFIIS to the RNA Pol II active site. Further genetic analyses in yeast supported this hypothesis (Hemming et al., 2000; Van Mullem et al., 2002).

We have shown here that the C-terminal domains of RPB-9 and TFIIS are conserved also in *C. elegans*. According to our study, RPB-9 is required to promote Integrator-mediated cleavage of the nascent transcript upon RNA Pol II backtracking at motif-dependent piRNA loci. Concomitantly, Beltran et al., (co-submitted) have reported a similar role for TFIIS. The fact that piRNA precursors are longer but can still be cleaved in *rpb-9* animals could perhaps indicate that TFIIS is still functional in these mutants, but not sufficient on its own to promote timely transcription termination. The same line of thought applies to *tfiis* mutants, which can terminate transcription, possibly thanks to a functional RPB-9 subunit, but do so in a defective manner (Beltran et al., co-submitted). Given these observations, we believe that it is possible that RPB-9 and TFIIS cooperate in the recruitment of Integrator activities to terminate transcription at piRNA loci. In the light of this, it would be interesting to explore, in the future, the precise molecular link between *rpb-9* and *tfiis* in *C. elegans*, and compare these results with the molecular data obtained in yeast. Overall, our study sheds new light on how the piRNA pathway utilises a core RNA Pol II subunit to guarantee high fidelity transcription.

## ACKNOWLEDGEMENTS

We thank Archana Yerra for critical reading of the manuscript, Charles Bradshaw and Peter Williams for helping with data submission. We thank the Gurdon Institute Media Kitchen for their support providing reagents and media. We thank Kay Harnish for his support managing the Gurdon Institute Sequencing Facility. We thank Julie Ahringer′s lab for kindly sharing RNAi bacterial clones. We are grateful for the Miska Laboratory members, especially Kin Man Suen, for helpful discussions and advice, and Marc Ridyard for laboratory management and maintenance of our nematode collection. We thank Toni Beltran and Peter Sarkies for exchanging data prior to publication.

## Funding

This work was supported by Cancer Research UK (C13474/A18583, C6946/A14492) and the Wellcome Trust (104640/Z/14/Z, 092096/Z/10/Z) to E.A.M; work in the Sarkies lab was funded by the Medical Research Council (Transgenerational Epigenetic Inheritance and Evolution) and an EMBO Young Investigator Award (to P.S.). A.C.B. was supported by a Marie Skłodowska-Curie Individual Fellowship (747666); G.F. was supported by an EMBO Long-Term fellowship, L.L. was supported by a Boehringer Ingelheim Fonds PhD fellowship. I.C.N. was supported by a Science without Borders Doctorate scholarship (205589/2014-6; CNPq, Brazilian Federal Government).

## AUTHOR CONTRIBUTIONS

Conceptualization, E.A.M.; Investigation, A.C.B., G.F., L.L., T.B., E.M.W., E.N., I.C.N., F.Br. and A.A.; Formal analysis, A.C.B., T.B., E.N., J.P. and F.Br.; Writing - Original Draft, G.F.; Writing - Review and Editing, E.A.M., A.C.B., L.L., A.A., P.S., T.B., F.Bu.; Visualization, L.L.; Supervision: E.A.M.; Funding Acquisition, E.A.M.

## DECLARATIONS OF INTEREST

The authors declare no competing interests. E.A.M. is a founder and Director of STORM Therapeutics Ltd. STORM Therapeutics had no role in the design of the study and collection, analysis, and interpretation of data as well as in writing the manuscript.

## SUPPLEMENTAL FIGURE LEGENDS

**Figure S1.**
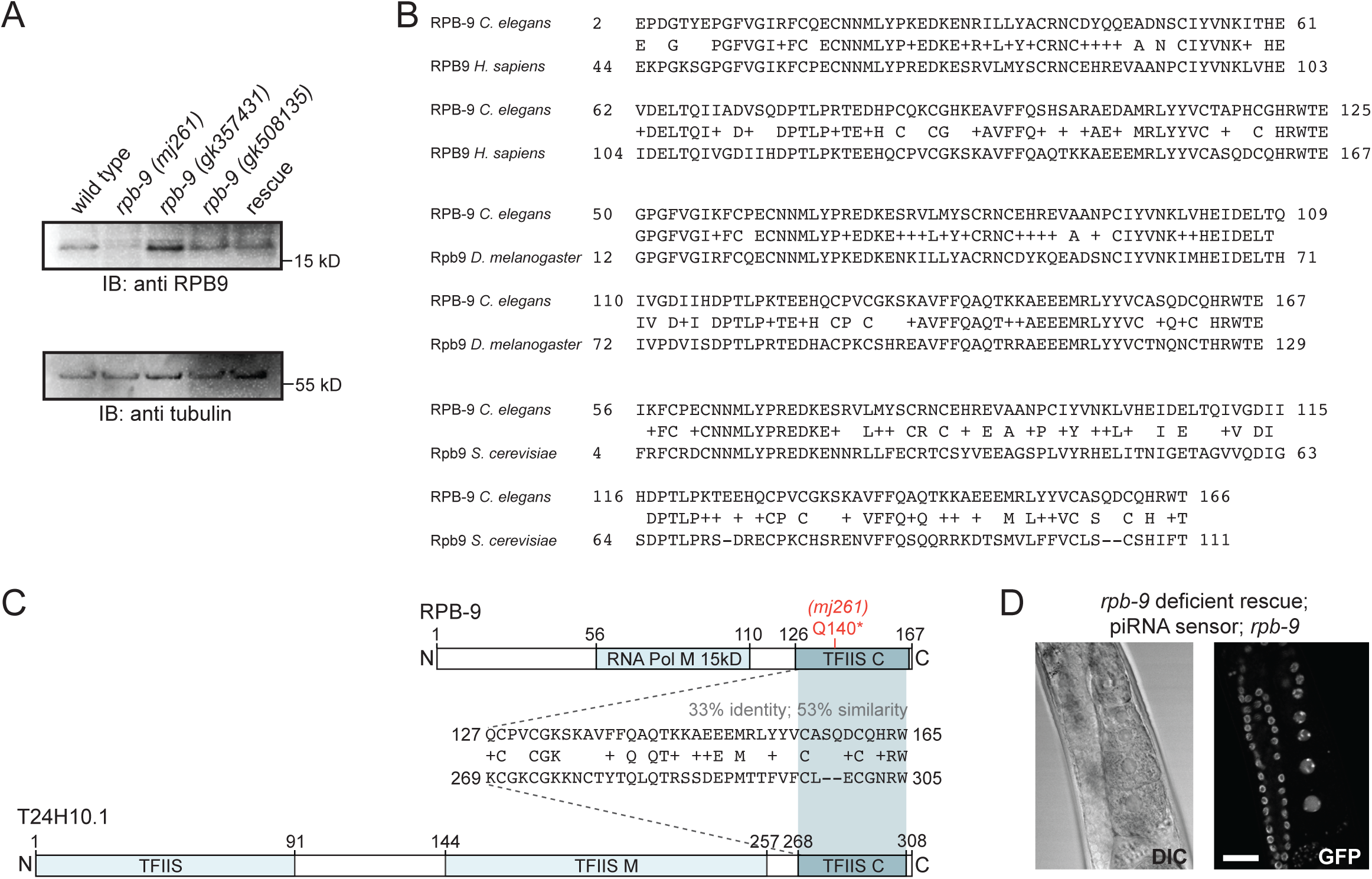
Related to Figure 1. **A**) Western blot quantification of RPB-9 expression in wild-type, *rpb-9 (mj261)* and *rpb-9* rescue animals. *rpb-9* alleles *gk508135* and *gk357431 (Thompson et al., 2013)* are additional controls that do not show piRNA sensor desilencing (data not shown). **B**) RPB-9 domain conservations. Alignments of the RPB-9 protein to its *H. sapiens, D. melanogaster* and *S. cerevisiae* homologs. Positive (NCBI BlastP) amino acids are indicated by ‘+′. **C**) Comparison of RPB-9 and the *C. elegans* TFIIS homolog T24H10.1 with alignment of the common TFIIS C domain. The introduced RPB-9 Q140* mutation is highlighted in red. Positive (NCBI BlastP) amino acids are indicated by ‘+′. **D)** Representative DIC and fluorescence microscopy images of piRNA sensor expression in deficient rescue *rpb-9 (mj261)*;*(mjSi89)* animals. Scale bar = 20 µm.

**Figure S2.**
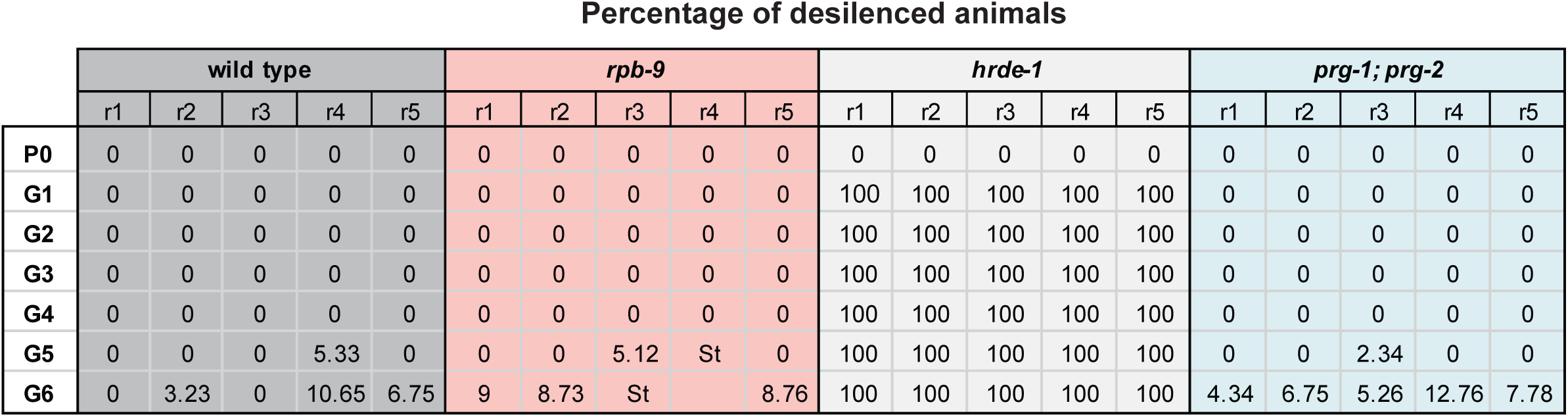
Related to Figure 2. **A**) Table of the inheritance of silencing experiment, showing the percentages of desilenced animals at each generation for individual replicates separately, for all the indicated genotypes.

**Figure S3.**
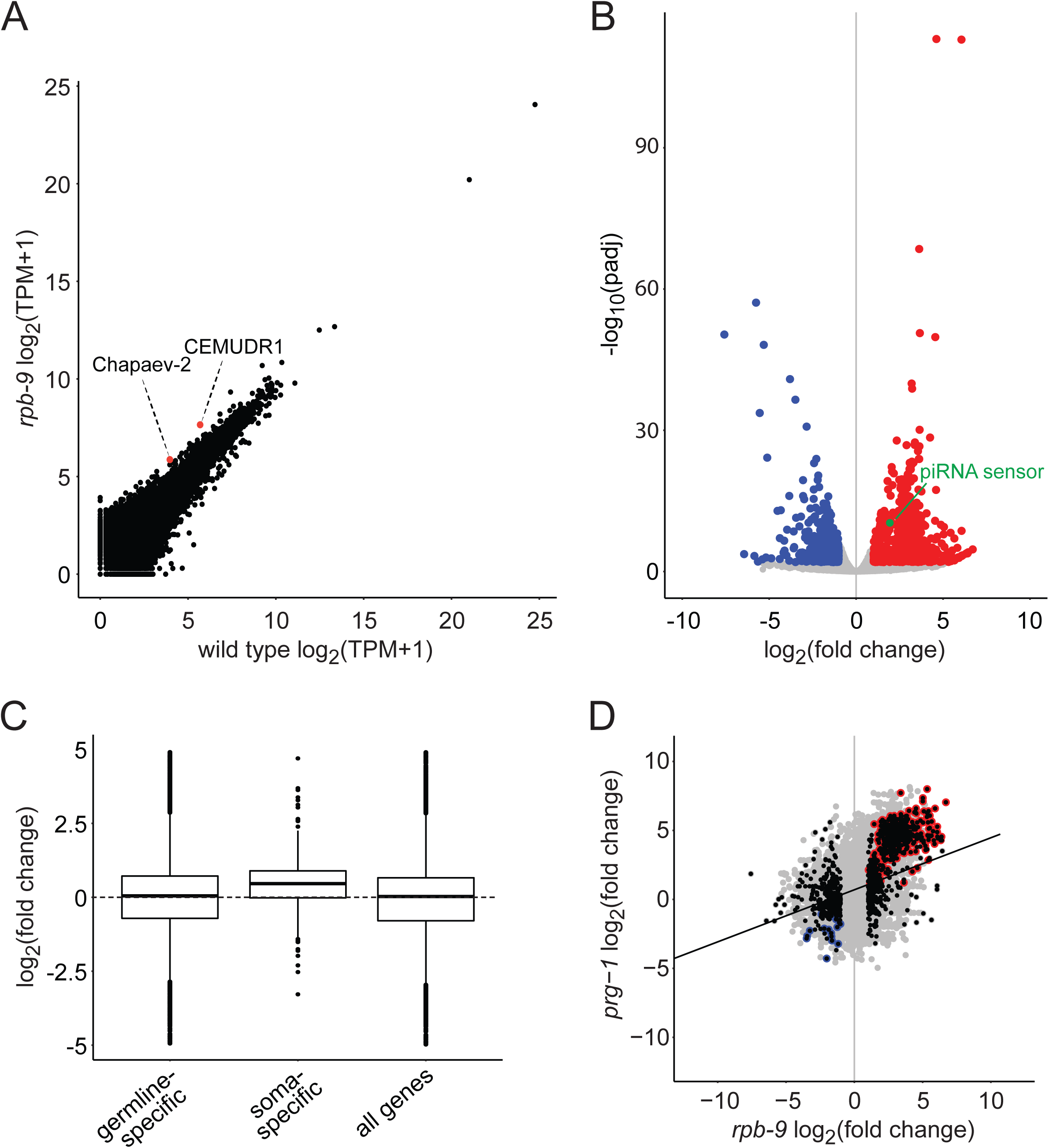
Related to Figure 3. **A**) Differential expression analysis of transposable elements in *rpb-9* (*mj261*) mutants versus wild type (total RNA “Ribo-Zero” RNA-seq libraries). **B**) Genome-wide differential expression analysis of *rpb-9* (*mj261*) mutants versus wild type (total RNA “Ribo-Zero” RNA-seq libraries). **C**) Differential expression analysis of germline-specific and soma-specific genes, as classified in (Reinke et al., 2004). All genes are shown for comparison. (Total RNA “Ribo-Zero” RNA-seq libraries). **D**) Pairwise correlation plots of *rpb-9 (mj261)* and gonad-dissected *prg-1 (n4357) (Reed et al., 2020)* transcriptomes. Differentially expressed genes in *rpb-9 (mj261)* are shown in black, shared upregulated genes in red, shared downregulated genes in blue. (Total RNA “Ribo-Zero” RNA-seq libraries).

**Figure S4.**
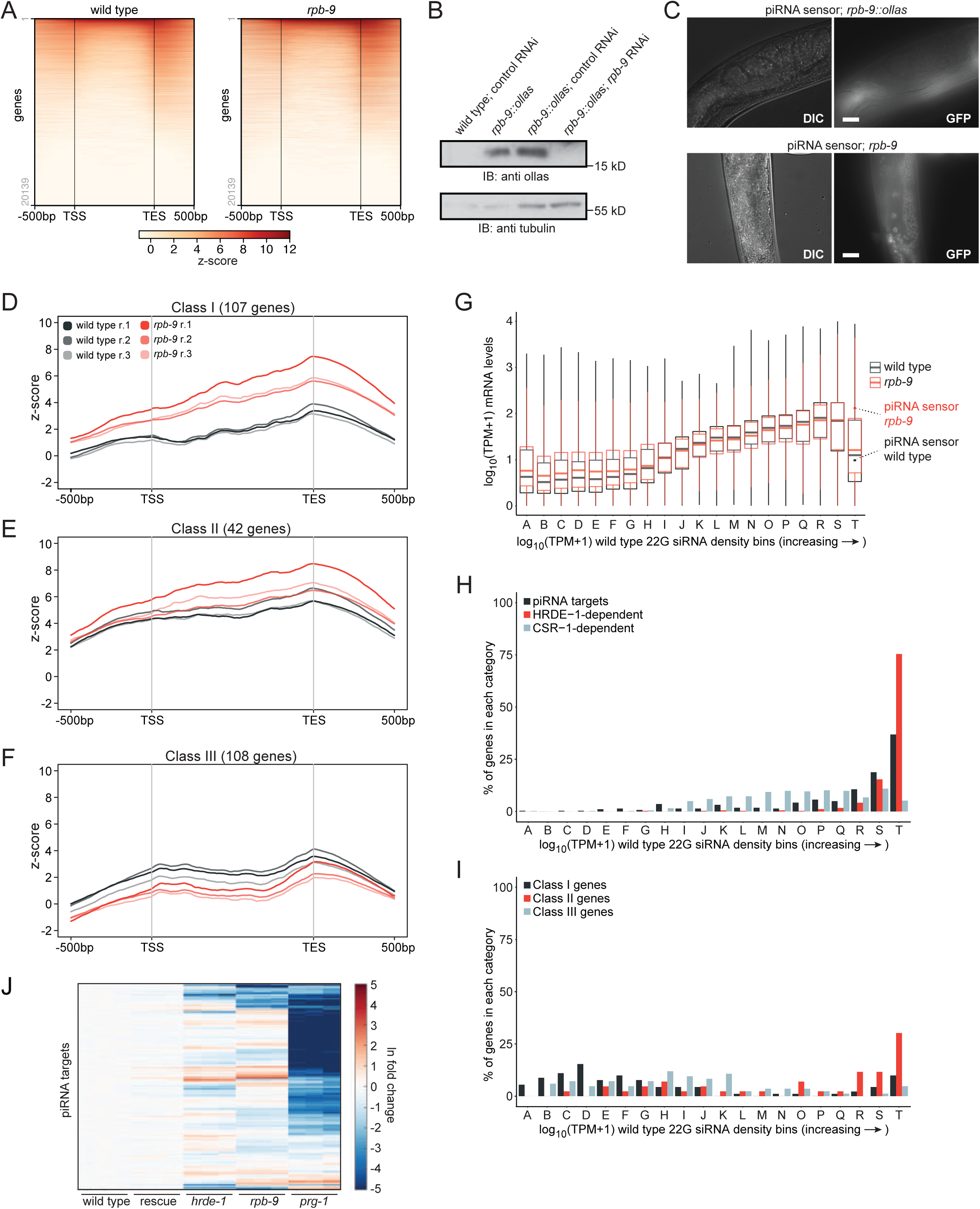
Related to Figure 4. **A**) Analysis of differential RNA Pol II binding (RPB-1/AMA-1 ChIP-seq) over all genes in *rpb-9 (mj261)* mutants compared to wild type. **B**) Western blot quantification of the amount of RPB-9 protein in the *rpb-9::ollas* line by western blot. *rpb-9* RNAi samples are shown for control. **C**) Representative DIC and fluorescence microscopy images of piRNA sensor expression in *rpb-9::ollas (mj604). rpb-9 (mj261)* is shown for control. Scale bar = 20 µm. **D-F**) Analysis of RNA Pol II binding at upregulated genes in *rpb-9 (mj261)* mutants, as defined in Figure S3B. Class I: upregulated genes with increased RNA Pol II binding (**D**); Class II: upregulated genes with invariant RNA Pol II binding (**E**); Class III: upregulated genes with reduced RNA Pol II binding (**F**). n=3 biological replicates are shown per each genotype. (RNA-sequencing filtering from total RNA “Ribo-Zero”RNA-seq libraries). **G**) Transcriptome binning according to increasing 22G siRNA density in wild-type animals (grey). Mean normalized *rpb-9 (mj261)* mRNA reads (red) are overlaid with mean normalized wild-type mRNA reads (grey). (total RNA Ribo-Zero RNA-seq libraries) The piRNA sensor transcript is highlighted in red (*rpb-9 (mj261)*) and grey (wild type). **H**) Distribution of piRNA targets (as defined in (Bagijn et al., 2012a)) and of HDRE-1- and CSR-1-dependent 22G siRNAs across bins as defined in G). **I**) Distribution of Class I, Class II and Class III genes (as defined in Figure S4D-F) across bins as defined in G). **J**) Cluster analysis of 22G siRNAs mapping to piRNA target genes in wild-type, *rpb-9 (mj261), rpb-9* rescue *(mjSi70), prg-1 (n4357)* and *hrde-1 (tm1200)* animals. (5’-independent small RNA libraries).

**Figure S5.**
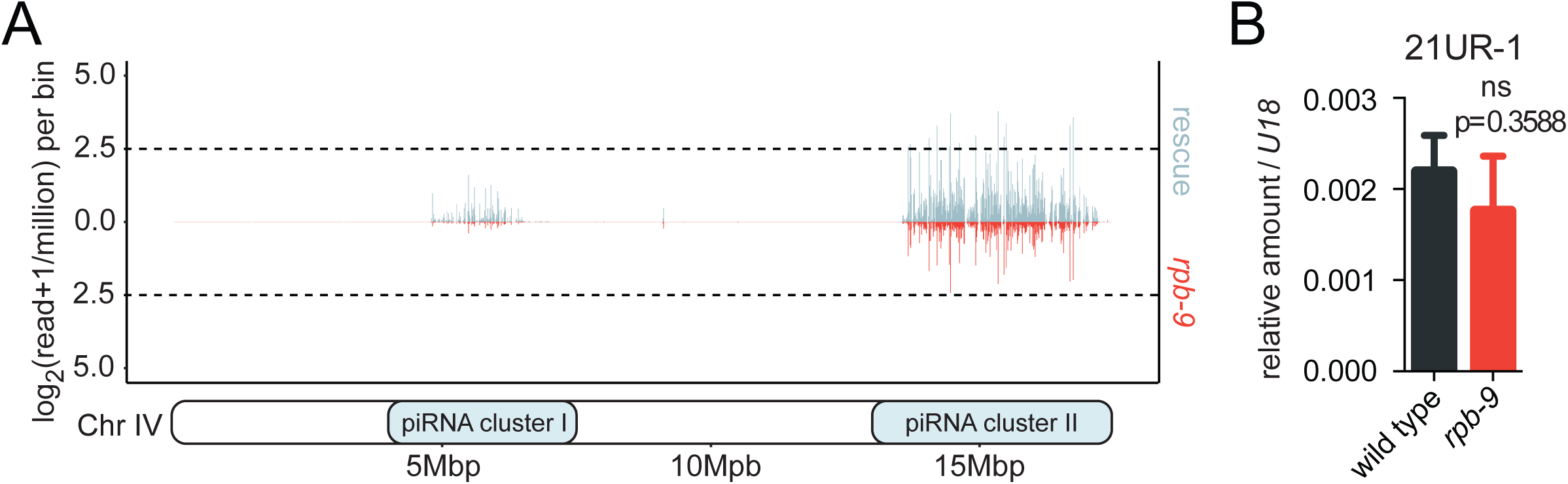
Related to Figure 5. **A**) Mature piRNA expression along chromosome IV coordinates (motif-dependent piRNA clusters I and II) in wild-type and *rpb-9* rescue animals. **B**) RT-qPCR quantification of mature piRNA 21UR-1 levels in wild-type and *rpb-9 (mj261)* animals. n=3 biological replicates, mean and SD shown (two-tailed t-test).

**Figure S6.**
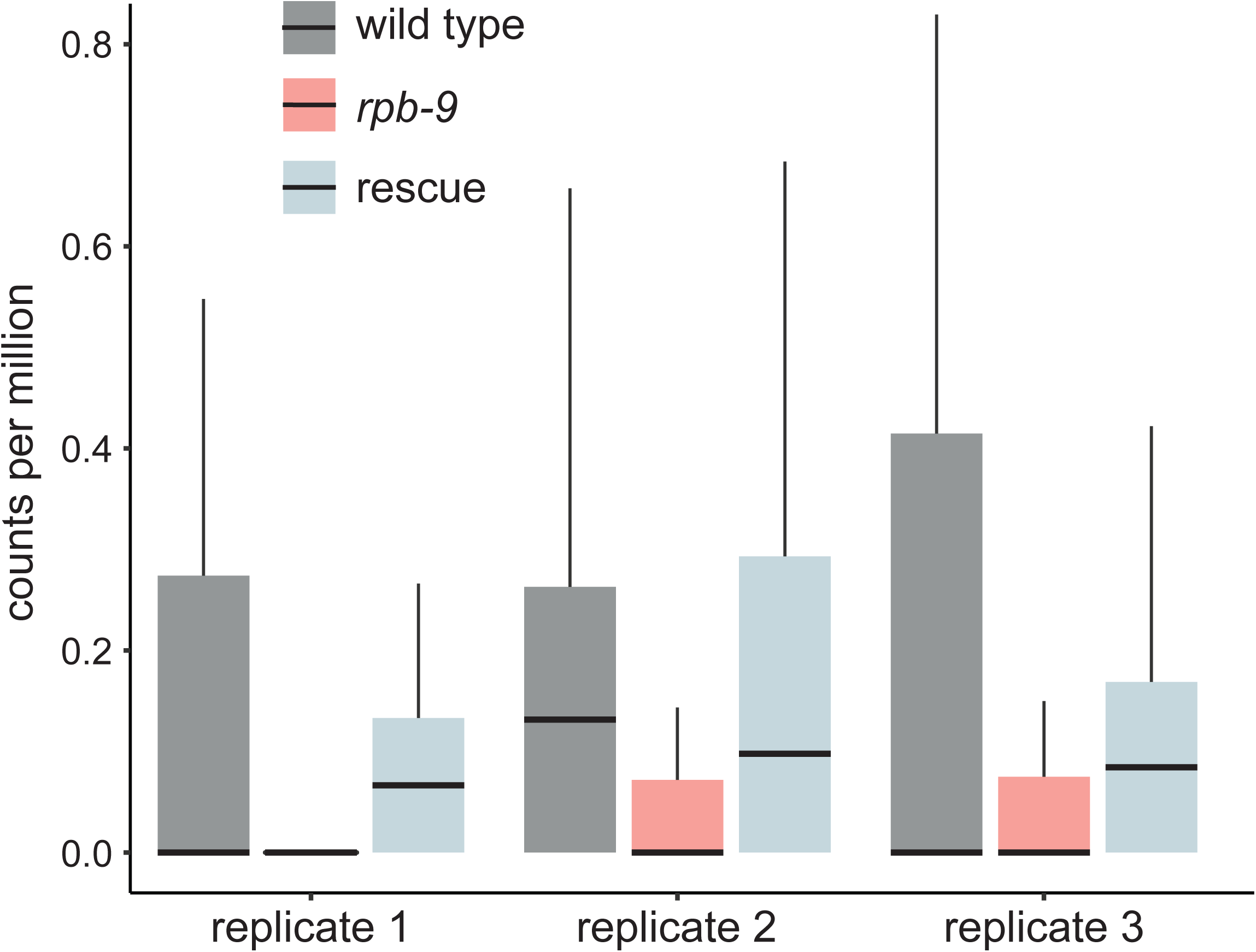
Related to Figure 6. Distributions of counts per million of 5′ monophosphate ends initiating in between +30 and +50 of piRNA TSSs, aggregated by piRNA locus.

## STAR METHODS

### RESOURCE AVAILABILITY

#### Lead Contact

Further information and requests for resources and reagents should be directed to and will be fulfilled by the Lead Contact, Eric Miska.

#### Materials Availability

Transgenic animals will be submitted to the *C.elegans* Genetics Center.

#### Data and Code Availability

RBP-9::ollas IP/MS data has been deposited to ProteomeXchange database with the accession number PXD018657.

All sequencing data generated in this study has been deposited to GEO with the accession number GSE149071.

### EXPERIMENTAL MODEL AND SUBJECT DETAILS

#### *C. elegans* culture

*C. elegans* were grown under standard conditions (Brenner, 1974) at 20°C using the *Escherichia coli* strain HB101 as a food source. Crosses were performed by mating males and females in a 5:1 ratio. All strains are listed in the Key Resources Table.

#### *C. elegans* transgenics - CRISPR injections

RPB-9 C-terminus was tagged with an Ollas peptide by injecting wild type N2 young adult animals with a mixture of target gene HR template (IDT oligos) (1 mg/ml), target gene CRISPR gRNA (Amersham Biosciences) (8 mg/ml), dpy-10 CRISPR gRNA (Amersham Biosciences) (2 mg/ml) as a marker, His-Cas9 (in-house bacterial purification) (5 mg/ml) in a buffer comprising 10 mM KCL and 10 mM Tris-HCL pH:8.0. Animals showing dumpy phenotype in the first generation after injection were screened for homozygous ollas tag insertion and backcrossed 3 times with the wild type N2 strain to get rid of CRISPR off target effects.

#### *C. elegans* transgenics - mosSCI injections

For targeted single copy transgene insertion, a mix of 20 ng/μl transgene construct, 50 ng/μl Mos transposase JL43 and 5 ng/μl for each co-injection marker (myo-2::mcherry::unc-54 and myo-3::mcherry::unc-54, or GFP markers as appropriate) was prepared and centrifuged for 30 min at 13,000 rpm (15,700 g) at 4°C. Microinjection pads consisted of 2% agarose flattened onto a cover slip. Prior to microinjections, a pad was moistened by gently exhaling onto it, then a drop of halocarbon oil 700 (Sigma Aldrich) was added to it. Young adult animals of the appropriate strain were transferred into the oil using an eyelash pick and flattened to the surface. Injections into gonad arms were performed using an Olympus IZ71 microscope equipped with an Eppendorf micromanipulator, Femtojet injection rig and Transfer-]Man joystick (Eppendorf). After injection, animals were left to recover in M9 medium, then transferred to individual plates and left to recover over-night at 20 °C. For selection of integrants, plates were placed at 25 °C and offspring were assessed for moving unc-119 rescue worms after 3 and again after 4 days. Once plates with motile worms had starved, they were chunked onto a new plate and any moving worms lacking the co-injection marker were singled onto individual plates. A maximum of ten individual animals was picked from each parental plate. Offspring were screened or re-individualised for absence of any Unc phenotype to generate homozygous lines. Transgene integration was validated by PCR of the insertion locus and transgene and, if applicable, by expression of a fluorescent protein. Details of the Mos system and selection process are described in (Frøkjær-Jensen et al., 2008; Frøkjæ-Jensen et al., 2012).

## METHOD DETAILS

### piRNA sensor EMS screen

After EMS treatment following standard protocols (Brenner, 1974), F2 or F3 offspring of mutagenized worms were sorted using a Copas Biosort large-particle sorter as described in (Bagijn et al., 2012b). Further details are as described previously (Ashe et al., 2012). Chromosome mapping and genotyping of mutations are described in (Weick et al., 2014).

### Cloning of mosSCI plasmids

Plasmid constructs were generated performing Multi-Site Gateway cloning (Invitrogen) according to manufacturer’s instructions. pDONR entry constructs were made either by amplifying the gene of interest including exons from genomic DNA, or by amplifying the spliced transcript from wild-type cDNA. All pDONR entry clones were confirmed by sequencing. pDEST vectors after LR reactions were confirmed by colony PCR on corresponding bacterial clones and by expression of transgene after injection into C. elegans. pDEST *C. elegans* expression constructs are detailed in (Frøkjæ-Jensen et al., 2012).

### piRNA sensor imaging

Representative single plan images were acquired on a Leica SP8 confocal microscope at 63X magnification. Immobilized live animals were used.

### Transgenerational memory assay

Three L4 larvae per genotype were plated on gfp RNAi-expressing bacteria (5 replicates) or empty vector L4440 bacteria (3 replicates). G1 worms were analyzed under a fluorescence microscope and one silenced worm per replicate per genotype was plated onto standard HB101 bacteria. At each generation, one silenced worm was singled from each plate to produce the next generation, and the remaining adult progeny was analyzed under a Kramer FBS10 fluorescence microscope. Worms were collected in M9, washed twice, quickly fixed in 70% ethanol and deposited onto a glass slide coated with a 2% agarose pad. At least 50 worms per replicate per genotype were counted at each generation. Germline nuclear GFP brightness was categorically scored as on or off.

Representative images were taken on a DM6B fluorescence microscope (Leica) with a motorized stage and a Leica DFC9000 GT CCD camera. Exposures: DIC 25msec, GFP 500msec. 40X magnification.

### RNA extraction and real-time quantitative PCR of spliced and unspliced mRNAs

Total RNA was extracted using Trizol reagent (AMBION-Life Technologies) and treated with Turbo DNAse kit (Invitrogen) according to the manufacturer’s instructions. 500ng of RNA were reverse-transcribed with random hexamers (Invitrogen) at 50°C for 1 hour using SuperScript III reverse-transcriptase (Thermo Fisher). Control reactions lacking enzymes were systematically run in parallel as negative controls. Real-time quantitative PCR were performed on 1ml of diluted RT reactions (1/5) using SYBR Green kit (Life Technologies) on a One Step Plus thermocycler (Thermo Fisher). All samples were run in duplicates. Expression levels were normalized to the reference gene *cdc-42*. Primer sequences are listed in the Key Resources Table.

### Cloning of the RPB-9 RNAi plasmid

The RPB-9 RNAi plasmid was constructed by cloning a 500bp fragment containing most of the RPB-9 cDNA sequence into the L4440 RNAi feeding vector (Timmons and Fire, 1998) via Gibson assembly following the manufacturer’s protocol. Briefly, 50ng of the *rpb-9* insert were added to 50ng of EcoRI-digested L4440 vector and incubated in Gibson master mix for one hour at 50°C. 1μl of the Gibson reaction was transformed into 50μl of Dh5α chemically-competent bacteria (NEB) and plated onto ampicillin-resistant plates. Resistant colonies were screened for correct insertions by Sanger sequencing with the M13 forward primer and an in-house reverse primer (GGCCTTTTGCTCACATGTTC).The following sequence was synthesized as g-Block (IDT): **GAGACCGGCAGATCTGATATCATCGATG**TATCTGCAATTAAAATCAAAGCTTGAAAATG TTTTATCATATTTTTTCAGAAAGATGAGCCAAGGGTATGATAATTACGATGATATGTACGATCAAAACGGTGCATCACCGGCGCCGAGTCAAAACGAAAAACCCGGGAAAAGTGGGCCTGGTTTTGTTGGAATCAAGTTTTGCCCAGAATGCAATAATATGCTGTACCCACGAGAGGATAAGGAATCACGAGTTTTGATGTATTCCTGCCGGAACTGTGAGCATCGTGAAGTCGCCGCTAACCCGTGTATCTATGTGAATAAGCTCGTTCACGAAATTGATGAGCTCACTCAAATCGTCGGAGATATTATTCACGATCCAACGCTCCCGAAGACTGAAGAACATCAATGTCCAGT CTGTGGCAAAAGTAAGGCTGTCTTCTTCCAGGCTCAAACAAAAAAGGCAGAAGAA**GAATTCGATATCAAGCTTATCGATACCGTC** (Bold: L4440 homology arms, which flank an EcoRI restriction site in the backbone L4440 vector).

### Western blotting

Western blotting of RPB-9 in the *rpb-9 (mj261)* and *rpb-9* rescue *(mjSi70)* (Figure S1A) was performed using a human anti-RPB-9 (NBP1-92344, Novus Biologicals) antibody and a mouse anti-tubulin (clone DM1A, SIGMA) antibody.

Western blotting of RPB-9 in the *rpb-9::ollas* line (Figure S4B) was performed using a mouse anti-tubulin (clone DM1A, SIGMA) antibody and a rat anti-ollas (NBP1-06713, Novus Biologicals) antibody.

Protein samples were prepared by boiling approx. 50 adult animals per genotype in NuPAGE™ (Thermo Fisher Scientific) sample buffer according to the manufacturer’s instructions. Denatured proteins were resolved on NuPAGE™ 4-12% BisTris gradient gels (Thermo Fisher Scientific) and wet transferred on 0.45mm pore-sized nitrocellulose membranes (Thermo Fisher Scientific) at 240mAmp for 2 hours. After over-night incubation with primary antibodies at 4°C, the membranes were incubated with HRP-conjugated secondary antibodies (GE Healthcare) for one hour at room temperature. Chemiluminescence Reagents (Pierce ECL Thermo Scientific or Immobilon Western Millipore) were applied to the membranes and the membranes were visualized on 18×24cm x-ray films in a dark-room.

### RPB-9::ollas Immunoprecipitation and Mass Spectrometry

N2 control or RPB-9::ollas animals were grown to young adult stage, washed 3X with M9 buffer and frozen in liquid nitrogen. Frozen worm balls were crushed with a metallic grinder and lysed with a buffer containing 25 mM Tris-HCL pH:7.5, 150 mM NaCL, 1.5 mM MgCl2, 0.1% Triton-X-100 and Complete Mini Protease inhibitor tablets (Roche, EDTA free). The lysate was sonicated using a BIORUPTOR with 10 cycle 30 SEC on/30 SEC OFF and then centrifuged for 30 min at 16,000 rcf (4°C) to remove insoluble material. BCA assay (Thermo Scientific) was used to determine total protein concentration of the supernatant. 4 mg total extract, 10 mg anti-ollas antibody (A01658-40, GeneScript) and 30 ml Protein A/G magnetic beads (Thermo Scientific) were used per technical replicate. In total, 4 technical replicates were used both for the N2 control and RPB-9::ollas. After 12 hours of incubation at 4°C in a rotating wheel, beads were washed 4 times with a buffer containing 25 mM Tris-HCL pH:7.5, 150 mM NaCL, 1.5 mM MgCl2 and Complete Mini Protease inhibitor tablets. Immuno-precipitates were then boiled with a 2X sample buffer for 15 minutes.

For the identification of RPB-9::ollas interactors, samples were separated on a 4%–12% NOVEX NuPage gradient SDS gel (Thermo) for 10 min at 180 V in 1x MES buffer (Thermo). Proteins were fixated and stained with coomassie G250 brilliant blue (Carl Roth). The gel lanes were cut, minced into pieces, and transferred to an Eppendorf tube. Gel pieces were destained with a 50% ethanol/50 mM ammoniumbicarbonate (ABC) solution. Proteins were reduced in 10 mM DTT (Sigma-Aldrich) for 1h at 56°C and then alkylated with 5mM iodoacetamide (Sigma-Aldrich) for 45 min at room temperature. Proteins were digested with trypsin (Sigma) overnight at 37°C. Peptides were extracted from the gel by two incubations with 30% ABC/acetonitrile and three subsequent incubations with pure acetonitrile. The acetonitrile was subsequently evaporated in a concentrator (Eppendorf) and loaded on StageTips (Rappsilber et al., 2007) for desalting and storage.

For mass spectrometric analysis, peptides were separated on a 20 cm self-packed column with 75-µm inner diameter filled with ReproSil-Pur 120 C_18_-AQ (Dr.Maisch GmbH) mounted to an EASY HPLC 1000 (Thermo Fisher) and sprayed online into an Q Exactive Plus mass spectrometer (Thermo Fisher). We used a 94 min gradient from 2% to 40% acetonitrile in 0.1% formic acid at a flow of 225 nL/min. The mass spectrometer was operated with a top 10 MS/MS data-dependent acquisition scheme per MS full scan. Mass spectrometry raw data were searched using the Andromeda search engine (Cox et al., 2011) integrated into MaxQuant suite 1.5.2.8 (Cox and Mann, 2008) using the Uniprot *C.elegans* database (August 2014; 27,814 entries). In both analyses, carbamidomethylation at cysteine was set as fixed modification while methionine oxidation and protein N-acetylation were considered as variable modifications. Match between run option was activated. Prior to bioinformatics analysis, reverse hits, proteins only identified by site, protein groups based on one unique peptide, and known contaminants were removed.

For the further bioinformatics analysis, the LFQ values were log2 transformed and the median across the replicates was calculated. This enrichment was plotted against the – log 10 transformed p-value (Welch t-test) using the ggplot2 package in the R environment.

### AMA-1 ChIP-seq

300,000 synchronised animals were grown to young adult stage on 140mm NGM plates and collected in an M9 buffer per replicate (3 for wild type and 3 for rpb-9 mutant). Animals were washed in the same buffer three times. Animals were frozen in liquid nitrogen and then grinded with a metallic mortar. These extracts were fixed with 1% formaldehyde at room temperature 9 min. The crosslinking was quenched with the addition of 0.125 M Glycine at RT for 5 min. for 4 min at 4000g (4°C). Following to two washes, a final wash with a buffer containing 150mM NaCl, 50mM HEPES/KOH, 1mM EDTA, 1% Triton-X 100, 0.1% sodium deoxycholate, protease and phosphatase inhibitors (Roche). Then the samples were sonicated for 30 cycles 30 SEC ON/30SEC OFF. Afterwards, the samples were centrifuged for 30 min at 13,000rpm (4°C). 1 mg of the supernatant was used for ChIP with 4µg of AMA-1 antibody (rabbit polyclonal, Novus, SDQ2357) incubated over-night rotating at 4°C together with 80µl magnetic protein A/G beads (Thermo Scientific). Samples were then washed twice in 150mM NaCl buffer, once with 500mM NaCl buffer, once with 1M NaCl buffer, once with TEL buffer (0.25M LiCl, 1% NP-40, 1% sodium deoxycholate, 1mM EDTA, 10mM Tris-HCl pH 8 and freshly added protease inhibitor) and finally twice in TE buffer (pH 8). Beads were then eluted twice in 60µl ChIP Elution Buffer (1%SDS, 250mM NaCl in TE pH 8) at 65°C. Eluates of same samples and input were treated with 2µl RNAse at 37°C for 1 hour, then with 1µl Proteinase K at 65°C to de-crosslink over-night.

### Library Preparation and Sequencing

ChIP-seq and polyA selected total RNA libraries were prepared with NEBNext® Ultra II DNA Library Prep Kit for Illumina. Optimum PCR cycles were determined via StepOnePlus qPCR using POWERTRACK SYBR Green reagents (Thermo Scientific). Ribo-zero selected total RNA libraries were prepared with NEBNext® Ultra II Directional DNA Library Prep Kit for Illumina.

ChIP-seq, 5’ dependent and independent small RNA and polyA selected total RNA libraries were sequenced on a Hiseq 1500 machine at the Gurdon Institute.

Total RNA was extracted using Trizol reagent (AMBION-Life Technologies) and treated with Turbo DNAse kit (Invitrogen) according to the manufacturer’s instructions. 1 μg total RNA was used for library preparation. Ribo-zero selected total RNA libraries were sequenced on a Hiseq 2500 at the Cancer Institute, Cambridge. All sequencing was performed with SE50. Short capped RNA sequencing from nucleoplasmic and chromatin gradients were sequenced at the LMS Sequencing Facility on a Hiseq 2500 with SE75.

### RNA extraction and real-time quantitative PCR of mature piRNAs

Total RNA was extracted using Trizol reagent (AMBION-Life Technologies) and treated with Turbo DNAse kit (Invitrogen) according to the manufacturer’s instructions. 5 μg of total RNA were oxidized (25mM NaIO_4_/1X Borate buffer) for 10 minutes at 25°C in the dark. 10ng of total RNA were reverse-transcribed with a TaqMan Small RNA Assays kit (Thermo Fisher) containing a gene-specific RT primer and a TaqMan MicroRNA Reverse Transcription kit (Thermo Fisher), according to the manufacturer’s instructions. Real-time quantitative PCR were performed on 1ml of RT reactions using TaqMan Universal Mastermix No AmpeErase UNG (Life Technologies) on a One Step Plus thermocycler (Thermo Fisher). All samples were run in triplicates. Expression levels were normalized to the reference gene U18.

### piRNA precursor length analysis

The analysis of piRNA precursor length was carried out as described in (Beltran et. al., 2020, cosubmitted), except for a bootstrap sample size of 5000 sequences owing to a greater number of detected piRNA precursor sequences across conditions. Degradation fragment analysis was carried out as described in (Beltran et. al., 2020, cosubmitted).

### RNA-sequencing data analysis

polyA and ribo-zero selected total RNA-seq libraries were mapped using STAR v2.5.4b. to *C. elegans* ce11 genome carrying the piRNA Sensor as an extra chromosome. STAR mapping parameters were “--outMultimapperOrder Random --outFilterMultimapNmax 5000 --outFilterMismatchNmax 2 --winAnchorMultimapNmax 10000 --alignIntronMax 1 --alignEndsType EndToEnd”.

Adapter sequences were trimmed from 5’dependent and 5’independent small RNA libraries with cutadapt v 1.15 (Martin, 2011) using “-m 14 -M 32 -a TGGAATTCTCGGGTGCCAAGG” parameters and were mapped against ce11 genome with the piRNA Sensor using STAR with ““--outMultimapperOrder Random --outFilterMultimapNmax 500 --outFilterMismatchNmax 1 --winAnchorMultimapNmax 10000 --alignIntronMax 1 --alignEndsType EndToEnd” parameters. Aligned RNA-seq reads were sorted and indexed with samtools v1.6 (Li et al., 2009). Total RNA read counts against the annotations were generated by featureCounts v1.6 (Liao et al., 2014) with parameters “-T 12 -M –fraction -t exon” from a GTF file.

22 nt small RNA reads with a 5’ G bias were extracted from 5’independent small RNA sorted bam files with a custom PERL script. Read counts with an anti-sense orientation to the transcriptional direction of annotated genes were extracted by featureCounts v1.6 using “-T 12 -M --fraction -s 2 -t exon” parameters. We next calculated the anti-sense 22G small RNA density over 20 equal bins by TPM normalisation for annotated genes. We then plotted corresponding mRNA TPM values in each bin and calculated the overlap between each bin with known HRDE-1, CSR-1(Zhang et al., 2011) and piRNA targets (Bagijn et al., 2012a). Anti-sense 22G small RNA profiles on the piRNA Sensor as well as transposons were normalized as reads per million (RPM). A GTF file containing annotations for transposable elements was generated by RepeatMasker (Smit et al., 2015) v 4.1.0 run with rmblastn version 2.2.27+ using RepeatMasker database version 20140131, against the ce11 genome. Read counts of total and small RNA libraries on individual transposons were calculated using uniquely mapped reads. Normalised counts, variance-stabilised counts, log2 fold changes and adjusted *P values* were obtained using DEseq2 v 1.18.1 (Love et al., 2014). For differential gene and transposon expression, candidates with a log2 Fold Change bigger than 1 or smaller than -1 as well as an adjusted *P value* smaller than 0.01 were considered as statistically significant differential expression. To overcome false-positive transposon differential expression, statistically significant upregulated transposon list was further filtered with the ones that span or overlap with the exons of statistically significant upregulated protein-coding genes.

piRNA abundance was analysed using 5’dependent libraries. Small RNA read counts for piRNAs were obtained using featureCounts v1.6 with “-T 12 -M --fraction” parameter and normalised as reads per million (RPM). Cluster analysis of 5’-independent small RNA libraries showing the fold change of small RNAs mapped piRNA targets in the indicated mutants compared to wild type. Fold change is displayed in natural log.

To compare changes in small RNA targeting piRNA targets in between worm strains (Bagijn et al., 2012a), small RNA reads per gene were counted and abundance calculated by correcting for library size using unique mapping reads (cutoff > 50 reads per million). The mean smallRNA abundance per gene was calculated, next the fold-change was calculated by dividing the mean abundance in mutant animals by the mean abundance in wild-type animals.

### ChIP-sequencing analysis

ChIP-seq libraries were mapped to *C. elegans* ce11 genome with the piRNA Sensor as an extra chromosome by BWA allowing multi mappers. sam files were coverted to bam format and then bam files were sorted and indexed using samtools v1.6. (Li et.al., 2009) ChIP-seq peaks were generated using MACS2 v 1.4.3 (Zhang et al., 2008) peakcall function against input libraries. Common peaks between individual replicates with a *P value* smaller than 0.01 were considered as significant peaks. Linear fold enrichment was calculated with MACS2 bdgcmp function by disabling --SPMR option per replicate. Log2 and Z-score transformations were performed on RStudio with a custom script. All individual replicates were combined with RStudio Rtracklayer and Genomic Ranges packages. Seqplots was used to generate density plots and heatmaps (Stempor and Ahringer, 2016). Class I, II and III genes were classified by mean log2 Fold Enrichment of combined replicates over gene bodies of statistically significant upregulated genes. Briefly, we considered genes for which mutant log2 Fold Enrichment was 1.5 times greater than that of wild type, as Class I. Similarly, genes with a mutant log2 Fold Enrichment 1.5 times smaller than that of wild type was considered as Class III. The rest of the upregulated genes remaining between these ratios were considered as Class II.

## QUANTIFICATION AND STATISTICAL ANALYSIS

Quantification and statistical analysis of polyA selected and ribo-zero depleted total RNA-seq, 5’dependent and independent small RNA-seq, AMA-1 ChIP-seq and piRNA precursor short RNA-seq is explained in methods. Adjusted p values of differential gene and transposon expression were calculated with DEseq2 v 1.18.1, which uses a negative binomial distribution. Small RNA counts were normalized as reads per million (RPM). Statistical significance for mature piRNA abundance and IP/MS were determined by two sample t test and Welch t test, respectively. To prevent zero values during log2 and log10 transformations, pseudo-count 1 was added for TPM and RPM normalisations.

## Notes

### Competing Interest Statement

The authors have declared no competing interest.

